# G2 stem cells orchestrate time-directed, long-range coordination of calcium signaling during skin epidermal regeneration

**DOI:** 10.1101/2021.10.12.464066

**Authors:** Jessica L Moore, Feng Gao, Catherine Matte-Martone, Shuangshuang Du, Elizabeth Lathrop, Smirthy Ganesan, Lin Shao, Dhananjay Bhaskar, Andy Cox, Caroline Hendry, Bastian Rieck, Smita Krishnaswamy, Valentina Greco

## Abstract

Skin epidermal homeostasis is maintained via constant regeneration by stem cells, which must communicate to balance their self-renewal and differentiation. A key molecular pathway, Ca^2+^ signaling has been implicated as a signal integrator in developing and wounded epithelial tissues[1, 2, 3, 4]. Yet how stem cells carry out this signaling across a regenerative tissue remains unknown due to significant challenges in studying signaling dynamics in live mice, limiting our understanding of the mechanisms of stem cell communication during homeostasis. To interpret high dimensional signals that have complex spatial and temporal patterns, we combined optimized imaging of Ca^2+^ signaling in thousands of epidermal stem cells in living mice with a new machine learning tool, Geometric Scattering Trajectory Homology (GSTH). Using a combination of signal processing, data geometry, and topology, GSTH captures patterns of signaling at multiple scales, either between direct or distant stem cell neighbors. Here we show that epidermal stem cells display dynamic intercellular Ca^2+^ signaling among neighborhoods of up to 10 cells that is surprisingly coordinated and directed through time across a pool of thousands of stem cells. We find that this collective coordination is an emergent property of the stem cell compartment, distinct from excitatory quiescent neuronal tissues. We demonstrate that cycling stem cells, specifically G2 cells, govern homeostatic patterns of Ca^2+^ signaling. Stem cells in different cell cycle stages dynamically regulate localization of the gap junction component Connexin43 (Cx43). Lastly, we uncouple global from local communication and identify Cx43 as the molecular mediator necessary for connectivity between local signaling neighborhoods. This work provides resolution in how stem cells at different stages of the cell cycle communicate and how that diversity of phases is essential for tissue wide communication and signaling flow during epidermal regeneration. Our approach provides a framework to investigate stem cell populations and their signaling dynamics, previously not possible.

## Introduction

Each day our bodies make and lose billions of cells[5, 6, 7]. This regenerative capacity is based on the ability to orchestrate fate decisions within an actively cycling stem cell pool, resulting in a balanced production of new stem cells (by division) and loss of cells (by differentiation or apoptosis). In epithelial regeneration across a number of organisms, these stem cell behaviors are directly coupled within local neighborhoods[8, 9]. Yet how stem cells communicate with their neighbors remains a largely unexplored area, due to the complexity of capturing signaling dynamics across space and time in a live, uninjured setting.

In response to injury and during development, epithelial stem cells are known to coordinate their regenerative behaviors via the Ca^2+^ signaling pathway[1, 2, 3, 4]. Implicated in a diversity of cellular functions, Ca^2+^ signaling is beginning to be understood as critical to stem cell function across systems[10, 11]. An abundance of *in vitro* studies have established that the temporal dynamics of Ca^2+^ signaling are tightly regulated and can differentially encode function, allowing for its versatility as a signaling pathway[12, 13, 14]. In the quiescent neuronal system, researchers have achieved sophisticated analyses of Ca^2+^ signaling across broad spatial and temporal scales in resting and stimulated neurons *in vivo*. However, in the regenerative stem cell context, the *in vivo* spatiotemporal characteristics of Ca^2+^ signaling and the fundamental mechanisms regulating these dynamics are still unclear.

Here, we set out to study how stem cells communicate with one another in regenerative contexts *in vivo*, focusing on the basal stem cell layer of the mouse epidermis. The basal layer is comprised of a heterogeneous pool of stem cells in various cell cycle stages. These cells balance self-renewal and differentiation behaviors to maintain homeostasis over the lifetime of the animal. Recent work has provided evidence of local coordination of these behaviors, where cell fates are influenced by their direct neighbors[8], but also of large-scale organization, where dynamic behaviours of diverse cell types are coordinated[15, 16]. In this highly regenerative tissue, communication between stem cells is necessary to carry out this coordination at different scales. While upregulation of intracellular Ca^2+^ levels has long been known to be essential for the terminal steps of epidermal differentiation in the skin[17, 18, 19, 20, 21, 22, 23, 24, 25], Ca^2+^ signaling dynamics and their regulation have not been explored in the basal stem cell layer. To investigate Ca^2+^ signaling activity *in vivo* in mammalian epidermal stem cells, we evolved our two-photon microscopy system[26, 27] to capture higher resolution images in Ca^2+^-sensor mice[28], allowing us to simultaneously capture dynamic Ca^2+^ signaling at the single cell level across thousands of cycling stem cells (**Figure 1A**).

**Figure 1:**
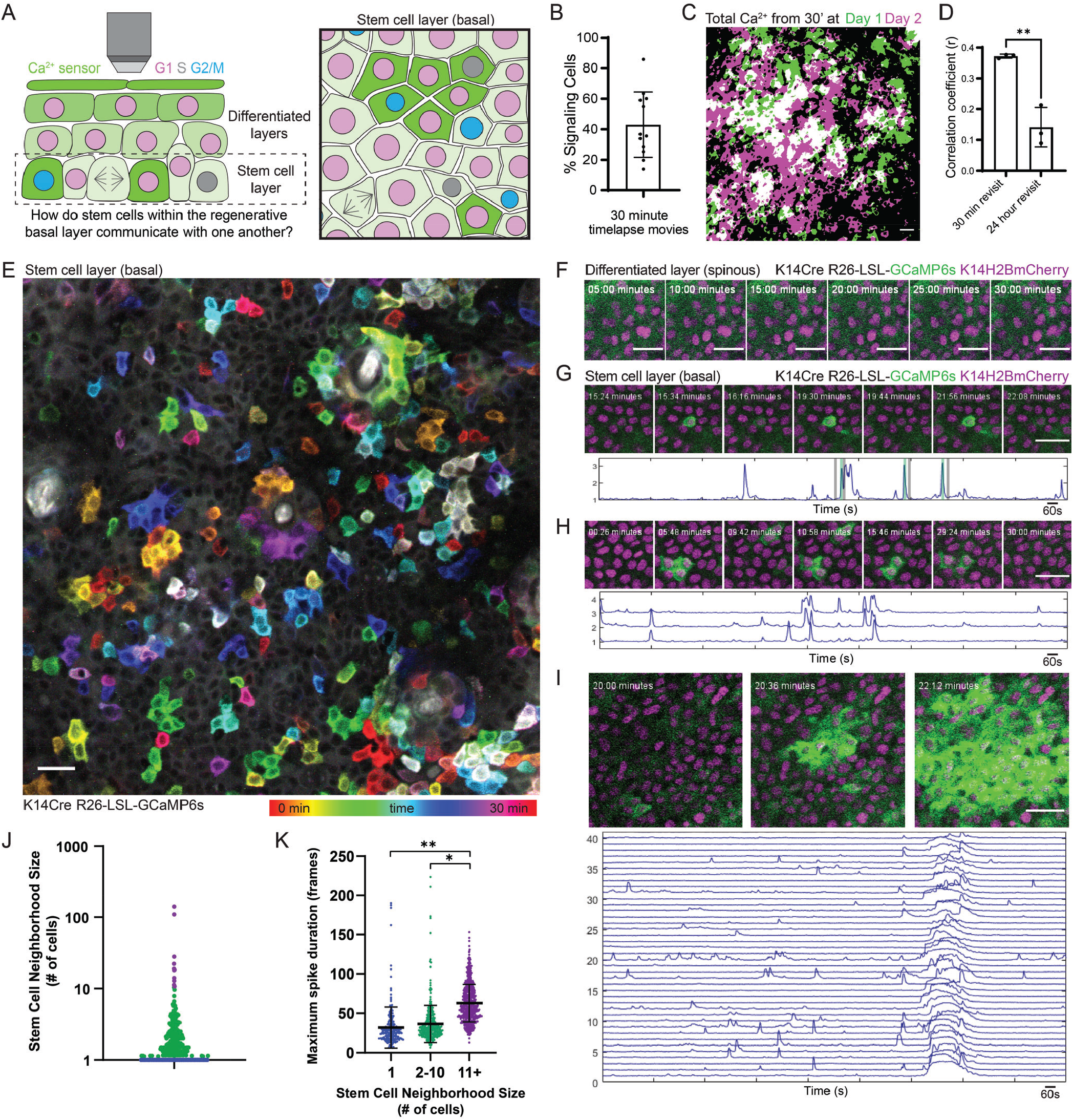
Epidermal stem cells carry out Ca^2+^ signaling across dynamic, local neighborhoods. **(A)** Schematic of intravital imaging of Ca^2+^-sensor mice, with a focus on the basal stem cell pool comprised of cycling cells. Cell cycle state is represented by nuclear color. **(B)** Percent of epidermal basal cells spiking at least once during 30-minute recording of Ca^2+^ signaling in a live mouse (K14-Cre; Rosa26-CAG-LSL- GCaMP6s; K14-H2BmCherry). N = 12 thirty-minute movies from 6 mice. **(C)** Composite image of the max intensity projections of all optical sections of a 30-minute timelapse at 0- (green) and 24-hours (magenta) of the same region of the epidermis. White indicates overlapping regions of Ca^2+^ signaling. Scale bars: 25 μm. **(D)** Correlation coefficient quantification of pixel overlap of Ca^2+^ signaling during 30-minute timelapses from revisits of the basal layer taken at 30-minute and 24-hour timepoints. ** = p *<*0.01, Student’s t test. N = 3 mice. **(E)** 30-minute timelapse video of the epidermal basal layer showing a diversity of spatiotemporal signaling patterns. Color scale represents time. Scale bars: 25 μm. **(F)** Lack of Ca^2+^ signaling in the differentiated spinous layer over 30 minutes of imaging (K14-Cre; Rosa26-CAG-LSL-GCaMP6s; K14-H2BmCherry). **(G)** Region of the basal stem cell layer where a single cell spikes repeatedly over 30 minutes of imaging (K14-Cre; Rosa26-CAG-LSL-GCaMP6s; K14-H2BmCherry). Normalized fluorescence intensity plotted over the duration of 30-minute timelapse is below. Black and green bars indicate timepoints corresponding to the snapshots above. Scale bars: 25 μm. **(H)** Different region of the stem cell layer, where a cluster of three cells spike repeatedly over 30 minutes of imaging. Normalized fluorescence intensity plotted over the 30-minute timelapse for each spiking cell is below. Scale bars: 25 μm. **(I)** Different region of the stem cell layer, where a large group of cells exhibit an intercellular Ca^2+^ wave (ICW). Normalized fluorescence intensity plotted over the 30-minute timelapse for 40 of the cells involved in the ICW is below. Scale bars: 25 μm. **(J)** Neighborhood sizes of cells with spatiotemporally localized Ca^2+^ signaling from 30-minute timelapse videos of the epidermal basal stem cell layer. Purple, blue, and green dots represent the three different spatial patterns of Ca^2+^ signaling. N = 6 thirty-minute timelapse movies from 3 mice. **(K)** Maximal spike duration (maximal number of frames between the start and end of individual Ca^2+^ events) for three different spatial patterns of Ca^2+^ signaling. * P = 0.0213, ** P = 0.0056, Nested One-way ANOVA, N = 6 thirty-minute timelapse movies.

Even with the capability of acquiring *in vivo* data, the representation and quantitative analysis of complex signaling patterns remains a significant challenge, due to both difficulties in visualizing and interpreting the global patterns within the signaling dynamics, as well as the cellular complexity inherent to regenerative tissues (i.e., thousands of stem cells, heterogeneous component cellular states, etc.). Therefore, analysis that properly captures the complexity of *in vivo* dynamics for any signaling pathway in mammals has been daunting and under-explored. Computational pipelines developed for signaling pattern analysis often use principal component analysis to identify cell assemblies (i.e., clusters of cells with similar dynamics) or rely on manual detection of signaling patterns[2, 29, 30], without addressing the relationship between constituent cells. More recent work has begun to analyze actively signaling cells as components of a signaling network, however these methods rely solely on correlation of signaling activity[31], missing related activity that could be slightly lagged in time.

To address this gap, we developed an unsupervised, data-driven computational analysis framework, named Geometric Scattering Trajectory Homology (GSTH)[32], to analyze and model signaling dynamics in stem cells for the first time. GSTH is based on a combination of graph signal processing (to capture signaling patterns over the tissue at various scales), data geometry (to map the structure of the data along time), and topology (to quantitatively characterize the data trajectory). GSTH facilitates the exploration of signaling patterns by quantifying within-sample dynamics and by allowing comparisons of global dynamics between experimental conditions. We demonstrate that this method can be widely applied and simultaneously overcomes many of the barriers inherent to the analysis of complex signaling dynamics, allowing for deeper understanding of stem cell signaling activity in many tissues.

These broadly applicable technological advances enable a bird’s eye view of tissue-wide signaling dynamics that was previously not possible. While coordination of intercellular signaling has typically been observed and predicted after external triggers, such as injury or mechanical stimulation, here we show that coordinated Ca^2+^ signaling is the baseline behavior in a homeostatic regenerative tissue. This elucidates how signaling networks can globally control tissue level principles. Here we find that cell cycle, G2 phase specifically, orchestrates these signaling dynamics, an unexpected reversal of our understanding of the relationship between signaling and cell cycle, with Ca^2+^ signaling usually implicated as a regulator of cell cycle progression. Further, we demonstrate that Cx43, a molecular player implicated in the past as an on/off switch for intercellular Ca^2+^ signaling, connects local signaling neighborhoods through global signaling coordination, underlying a large scale, homeostatic connectivity unobserved prior to this work. Together, our results provide insight into how a heterogeneous pool of stem cells comes together as a community to regulate the essential Ca^2+^ signaling pathway at unprecedented scale to maintain proper homeostasis.

## Results

### Epidermal stem cell pool displays local and dynamic neighborhoods of Ca^2+^ signaling

Under homeostatic conditions, epidermal stem cells either progress through the cell cycle towards division or exit into the suprabasal differentiated layer **(Figure 1A)**. Proper coordination of these behaviors is necessary for healthy skin regeneration and requires communication among stem cells. To understand the characteristics and regulation of stem cell communication, we turned to Ca^2+^ signaling, which is essential for regenerative behaviors, such as proliferation, in other epithelial contexts[1, 2, 3, 4]. We generated mice with a Ca^2+^-sensor expressed in all epidermal cells (K14-Cre; Rosa-CAG-LSL-GCaMP6s) and combined this with a nuclear marker (K14-H2BmCherry). Live imaging of the mouse basal stem cell layer[26, 27] revealed highly variable levels of participation from the basal cells, with 43.1 ± 21.4 % of cells showing at least one Ca^2+^ transient within a given 30 minute time frame (**Movie 1, Figure 1B**). To determine the characteristics of this communication, we asked whether homeostatic Ca^2+^ signaling is restricted to specific cells or shared across all cells within the basal stem cell layer. To this end, we quantified Ca^2+^ signaling in a large (500 μm by 500 μm) region, encompassing about 2,500 epidermal basal cells, over a period of 24 hours, during which many stem cells cycle through different cell cycle phases (**Figure A1A**). Comparison of active Ca^2+^ signaling across the 24 hour time period revealed that Ca^2+^ signaling is not spatially persistent (**Figure 1C, A1B**) but rather changes regionally with time. To quantify this, we measured the degree to which pixels with thresholded Ca^2+^-sensor fluorescence intensity overlapped at 0hr and 24hr. The correlation of signaling activity across 24 hours (0.1411 ± 0.06416) was significantly lower than the signaling correlation of same region just 30 minutes later (0.3723 ± 0.0064) (**Figure 1D, A1C**). Together, these results demonstrate how stem cell communication via intercellular Ca^2+^ signaling is dynamic across tissue domains and pervasive throughout the basal stem cell layer.

To understand how stem cells orchestrate Ca^2+^ dynamics on a shorter timescale within a field of connected epithelial stem cells, we again imaged large epidermal regions every 2 seconds for 30 minutes and temporally color coded each frame of the timelapse movie to simultaneously visualize all the Ca^2+^ signaling patterns (**Figure 1E**). We observed distinct spatiotemporal patterns of Ca^2+^ signaling within the stem cell layer: in some cases single cells spiked quickly in isolation, whereas in other cases neighborhoods of cells spiked simultaneously or in a propagating wave. The dynamic nature of these intercellular signaling events is a feature of the epidermal stem cell layer and does not characterize the directly above differentiated and quiescent suprabasal layer, which shows no signaling activity (**Figure 1F**).

To systematically quantify Ca^2+^ transients from each individual cell in the stem cell layer and to understand the relationship in time and space between these Ca^2+^ transients, we adapted existing tools[29, 33] to segment individual cells, developed a peak finding system, and defined a simple graph in which nodes were connected if the cells they represented were direct neighbors (within 1 μm of each other) and spiked within 10 seconds of their neighbor. We could then quantify the number of connected nodes or stem cells in each clustered signaling neighborhood and explore the temporal dynamics for each neighborhood size. Using this approach, we found that most events within the stem cell layer were either single cells that spiked in isolation from their neighboring cells (65.88 ± 2.65%; **Figure 1G, 1J**) or spatiotemporally clustered transients across 2 or more neighboring cells (31.27 ± 1.96%; **Figure 1H**). We also observed rare Ca^2+^ signaling waves that occurred across hundreds of cells (**Figure 1I**). We next quantified the duration of the longest Ca^2+^ transient per cell, which we termed maximal spike duration. We found that Ca^2+^ transients in larger neighborhoods of cells persist longer than in single cells or small neighborhoods (**Figure 1K**). These data show that the stem cell layer is characterized by local patterns of Ca^2+^ signaling across neighborhoods of mostly 1 to 10 cells with distinct temporal characteristics and that these regions of signaling change as the basal epithelium turns over.

### Geometric Scattering Trajectory Homology (GSTH) reveals long-range signaling across the stem cell pool, distinct from excitatory, quiescent systems

Epithelial cells have been shown to display coordinated signaling across a tissue in response to injury[20, 34, 35]. However, it’s unclear whether the local neighborhoods of Ca^2+^ signaling we observe during homeostasis occur at random or are coordinated with one another. Understanding signaling dynamics of any molecular pathway at a large scale *in vivo* represents a formidable challenge, especially in highly dynamic regenerative tissues. This is due to both the spatiotemporal heterogeneity of signaling dynamics as well as the complexity inherent to the tissue (i.e., thousands of stem cells, heterogeneous states of the signaling cells, dynamic cellular behaviors, etc.). To analyze signaling across multiple scales, we developed a method called GSTH - *Geometric Scattering Trajectory Homology*[32], which captures spatial and temporal patterns of signaling in a highly applicable manner and allows us to compare the Ca^2+^ signaling behavior of stem cells under many conditions.

To motivate GSTH, consider the problem of representing a signal on a set of cells (here we have epidermal stem cells arranged in planar spatial patterns). If we simply describe the signals as a vector of values on an indexing of cells, then we could not compare signaling patterns from different tissues, as specific cellular coordinates are not matched between tissues. Therefore, the signaling description has to be invariant to permutations in cell indexing, shifts in the signal, and even differences in the number of cells. To address this issue, in classic signal processing, researchers use frequency domain descriptions, such as the Fourier transform (FT), which describe the periodicity rather than the time- or space-specificity of signals. With the prevalence of graph-structured data, there is an emerging field of graph signal processing[36], in which researchers have invented the analogous graph Fourier transform (GFT)[37]. In our case, the graph consists of cells as vertices, and edges are determined by physical adjacency (**Figure 2A-Step 1**). However, the GFT (and the FT) is usually only suitable for describing signals with global periodic patterns. More localized signaling patterns can be described using wavelet transforms. Here, we use a graph wavelet transform[38], which can capture both localized and diffuse signaling patterns across the cellular graph.

**Figure 2:**
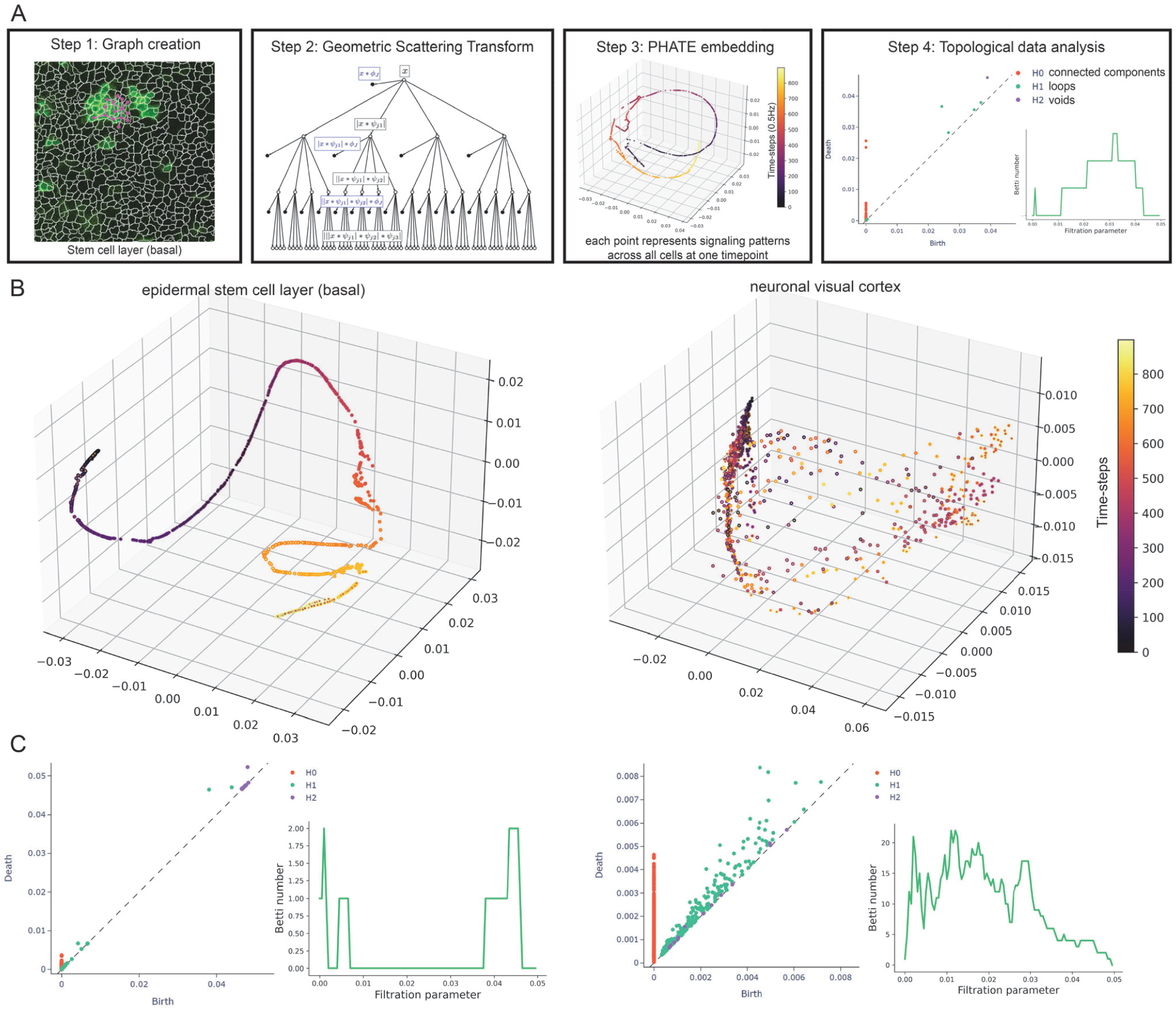
GSTH analysis reveals directed and coordinated Ca^2+^ signaling patterns across the basal epithelium. **(A)** GSTH workflow. Step 1: The cellular graph is created based on spatial adjacency (shown superimposed on a segmented image of basal stem cell layer Ca^2+^ signaling). Step 2: Timepoint embeddings are created using the geometric scattering transform. Step 3: PHATE (dimensionality reduction method) visualization of signaling time trajectory. Step 4: Topological data analysis via persistence diagrams of each trajectory (*H*0: connected components, *H*1: loops, *H*2: voids) and featurized representations of the diagram with Betti curves. **(B)** Representative PHATE visualization of coordinated Ca^2+^ signaling in the homeostatic epithelial stem cell layer (left) versus disorganized Ca^2+^ signaling in the quiescent neuronal visual cortex (right). **(C)** Corresponding persistence diagrams of Ca^2+^ signaling time trajectories in the homeostatic epithelial stem cell layer (left) versus the quiescent neuronal visual cortex (right). Each point corresponds to a topological feature in the trajectory, which appears at a certain birth time and disappears at a death time. As an example, green points represent *H*1 features that correspond to the formation of loops in the trajectory while purple dots represent *H*2 features that correspond to the formation of voids. The further they are from the respective diagonals, the longer they exist, i.e., the larger their persistence. To the right, examples of corresponding Betti curves of *H*1 loop features.

However, one scale of wavelet transforms is not sufficient to capture all the invariances we need in the signals. For instance, two signaling patterns, at two points in time, could be similar but just shifted by one cell, or similar overall but different cell to cell. To capture a broader notion of similarity between signals at different timepoints, we look at signaling at all scales of granularity. To achieve this, we do not simply use raw signals but also versions of these signals that are diffused to different scales. These diffusions are a natural part of a particular type of graph wavelet, called a *diffusion wavelet*[38]. GSTH uses multiple scales of diffusion wavelets, whose coefficients are locally averaged in a *geometric scattering transform* (**Figure 2A-Step 2**). Specifically, the geometric scattering transform uses statistical moments of wavelet coefficients over all cells to achieve permutation invariant descriptions of the signaling patterns, giving us a way of capturing the intrinsic signaling pattern unaffected by rotations and shifts of the signal on the tissue (see [39] for permutation invariance results).

Not only are our signals patterned in space, but they are also patterned in time. To understand signaling dynamics, GSTH uses a time series of geometric scattering transforms visualized in a dimensionality reduction visualization called PHATE[40]. PHATE reveals the emergent structure of the signaling time trajectory due to its ability to preserve manifold distances in low dimensions, whereas methods such as UMAP and tSNE tend to shatter the trajectory (as shown in Methods). Each point in the PHATE plot represents the global and local Ca^2+^ signaling pattern of thousands of cells during one time step (**Figure 2A-Step 3**). The trajectory is then colored by the index of time steps, so that time steps with similar Ca^2+^ signaling patterns are near each other in 3-D space. From these trajectories, we can then begin to understand how Ca^2+^ transients pass between cells in the stem cell pool in space and time.

Finally, to develop a quantitative descriptor of the entire spatiotemporal signaling pattern, we convert the PHATE time trajectory into a persistence diagram[41] (**Figure 2A-Step 4**). Such topological descriptors help quantify how the signaling patterns are connected or related across different scales of time. These descriptors use abstract topological features that appear in the trajectory as points are merged with one another using a coarse-graining operation. At each level of granularity, persistence diagrams quantify the shape features (holes, voids, and connected components) that emerge in the data. Moreover, persistence diagrams of dynamic trajectories from different mice can be readily compared via well-defined Wasserstein distances[42], which we employ here.

To capture spatial and temporal patterns of signaling across different scales in the basal stem cell layer, we applied GSTH to the 30 minute timelapses (900 time steps) of homeostatic Ca^2+^ signaling in this compartment. We first constructed a nearest-neighbors cellular graph, generated scattering coefficients for each timepoint, and then used PHATE to visualize the time trajectory. Each point in the time trajectory was color-coded based on timepoint and its position in space revealed the similarity or differences of the tissue’s Ca^2+^ signaling pattern from timepoint to timepoint. Our analyses revealed smooth time trajectories between timepoints, showing that Ca^2+^ transients steadily diffuse to stem cell neighbors in a directed and coordinated manner (**Figure 2B, A2A**). This analysis revealed an emergent property of this compartment, that localized signals are coordinated and patterned in time and space across larger tissue scales.

To determine how these PHATE trajectories from the regenerative epidermis compared to other tissues, we next applied GSTH to a classic example of Ca^2+^ signaling in the nervous system using previously published recordings of Ca^2+^ signaling from 10,000 neurons of the primary visual cortex[43, 44]. Spontaneous activity from the primary visual cortex has been shown to not be organized topographically during the resting state. Neurons can be connected via long processes, and so we used correlation between neurons’ Ca^2+^ signals to create a neuronal graph (instead of the nearest-neighbors graph built for epidermal cells). We then followed the same steps to generate scattering coefficients for each timepoint. Our analysis revealed markedly discontinuous, scattered time trajectories with PHATE, indicating less spatially and temporally coordinated signaling across the tissue and more abrupt changes in signaling patterns over time (**Figure 2B**).

To establish the generality of the GSTH method, we studied whether GSTH could differentiate between two different neuronal dynamics (which would represent a more subtle difference than comparisons between epidermal and neuronal signaling). We thus applied GSTH to an additional published visual cortex dataset[43, 44] (this time the mouse was stimulated with images) to see whether we would detect differences when compared with the spontaneous signaling in the visual cortex. While we observed some similarity in signaling patterns (in that they both were not as organized in time as the epidermal basal cells), the stimulated neuronal dataset displayed a much narrower, lower dimensional state space (locally some points with similar colors were near each other), contrasting with the dispersed PHATE plots from the unstimulated visual cortex datasets (**Figure A3A**). This demonstrated that the visual cortex shows less random patterns of Ca^2+^ signaling when stimulated. To investigate topological differences in signaling patterns of the stimulated and unstimulated visual cortex, we carried out the final step of GSTH, calculating persistence diagrams and Betti curves (**Figure A3B)**. We observed that the spontaneous neuronal signaling trajectory contained more *H*1 and *H*2 features, representing loops and voids in the trajectories. These loop and void features demonstrate that the topology of the spontaneous condition’s trajectory is more complex, scattered, and chaotic than the stimulated condition. This comparison demonstrates the applicability of GSTH to detect differences in global signaling patterns in a variety of systems.

To quantitatively compare the neuronal and epithelial stem cell layer datasets, we again carried out the final step of GSTH, visualizing persistence diagrams and Betti curves of *H*1 features (**Figure 2C**). If there are deviations from the main trajectories, then these will form persistence features that appear earlier because they create loops at low thresholds of point connection. By contrast, smooth trajectories only create large scale loops appearing later in the persistence diagram. The persistence diagrams were markedly different, with many features appearing and disappearing at all scales for the neuronal dataset, revealing a complex data geometry. By contrast, the persistence diagram of the epidermal stem cell layer had only a few noise features that quickly disappeared and then only large scale loops, with looping dynamics appearing much later in the persistence diagrams. Further, a prominent *H*2 feature or void (like the inside of a hollow ball) appeared at a late persistence stage in the epidermal dynamics, showing an area of the state space that was not entered in these dynamics. By contrast, the neuronal persistence diagram had several low-persistence *H*2 features that appeared and disappeared quickly, revealing more complex topological features in the neuronal dataset. Thus, we find that the epidermal stem cell pool orchestrates tissue-wide coordinated and directed Ca^2+^ signaling through time, demonstrating the spatial and temporal connectivity of information flow in the basal epithelium across multiple scales during homeostasis. This broadly applicable tool uncovered an unexpected emergent phenomenon of a regenerative stem cell compartment that is distinct from completely synchronized or random patterns of Ca^2+^ signaling described in other contexts.

### G2 cells are essential in mediating directed, coordinated patterns of signaling in the epidermal stem cell pool

We show that that the basal stem cell layer coordinates localized signaling events across the tissue to carry out long-range information flow. However, we fail to understand what the cellular mechanisms underlying this coordination are. The basal stem cell layer of the epidermis is a dynamic environment characterized by a heterogeneous distribution of cells in various phases of the cell cycle[15]. We wondered whether all basal cells (as heterogeneous cellular units) have the same roles towards tissue-wide, collective Ca^2+^ signaling or if cells in specific cell cycle stages regulate Ca^2+^ dynamics differently, as shown in other systems[11, 45, 46, 47]. To address this, we used the Ca^2+^-sensor combined with the Fucci cell cycle reporter that fluorescently labels G1 and S cells in red[48]. In our system, Fucci negative cells would be in G2 or mitosis, allowing us to look at two halves of the cell cycle and make comparisons. We observed clusters of Ca^2+^ transients propagating across cells of G1/S and G2/M cell cycle stages (**Figure 3A**). Quantification of the overall Ca^2+^ signaling activity revealed similar overall competencies to participate in signaling across all neighborhood sizes (**Figure A4B, A4C, A4D**). To next investigate the the spatiotemporal characteristics of Ca^2+^ signaling across cells of different cell cycle stages, we created embeddings of the cells (henceforth referred to as the cellular embeddings), based on all points in time, using the wavelet coefficients computed during GSTH. By concatenating these wavelet coefficients at every timepoint, we produced a cellular embedding for each cell that encompasses Ca^2+^ signaling information from that cell and its close neighbors across all timepoints. We finally applied PHATE to generate low-dimensional PHATE embeddings for each cell. We then colored each point in the PHATE plot based on its cell cycle stage to see G1/S versus G2/M cells in terms of their Ca^2+^ signaling patterns in 3-D space. We found that G2/M cells appeared to cluster together, showing related Ca^2+^ signaling patterns (**Figure 3B, A5A**). This was in contrast to G1/S cells, which were highly dispersed across the PHATE plots, indicating heterogeneous patterns of signaling. Ca^2+^-sensor fluorescence traces from the timelapses were consistent with these findings (**Figure 3C**). Collectively, these results demonstrate that G2/M cells display Ca^2+^ signaling patterns that are more similar to each other in spatial and temporal dimensions than G1 or S cells.

**Figure 3:**
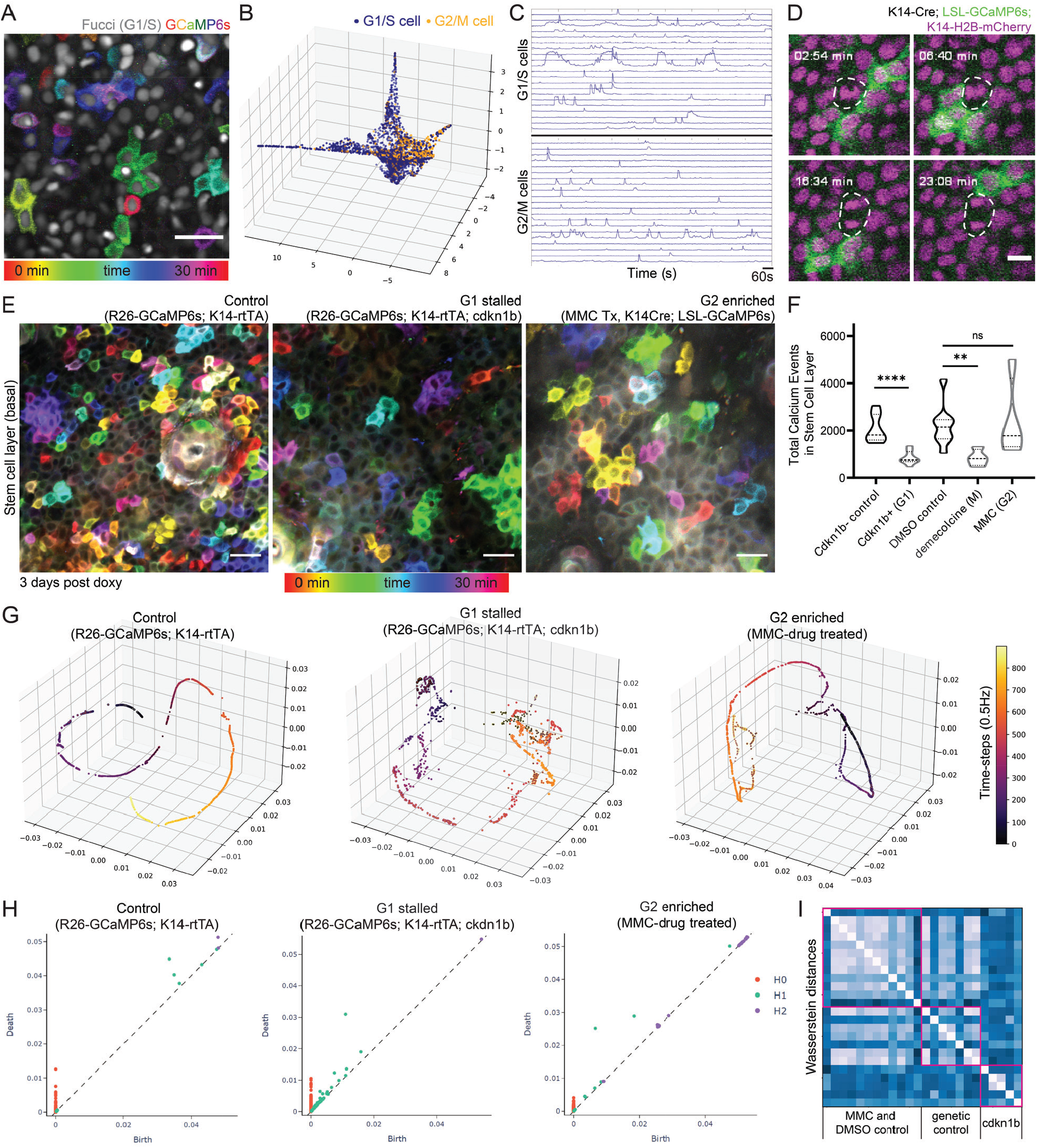
G2 cells are essential in mediating coordinated patterns of signaling. **(A)** Representative example of mitotic cell with lack of Ca^2+^ signaling in live mice (K14-Cre; Rosa26-CAG-LSL-GCaMP6s; K14-H2BmCherry). Nuclei in magenta and Ca^2+^ levels represented in green. Mitosing cells were identified via visual mitotic spindles in the nuclear signal. Scale bar: 10 μm. **(B)** Max intensity projection of 30-minute timelapse video of the epidermal basal layer of Ca^2+^-sensor mouse with a cell cycle reporter (K14-Cre; LSL-GCaMP6s; Rosa26p-Fucci2). Only the mCherry-hCdt1 expression is visible (grayscale). Time represented as color scale. Scale bar: 25 μm. **(C)** Representative PHATE plots of cellular embeddings. Each dot represents a single cell; its position in space represents similarity of its Ca^2+^ signaling to other cells’ signaling; each cell/node is colored by its cell cycle state based on Fucci signal. **(D)** Representative heatmap of Ca^2+^ activity over time in G1/S cells versus G2 cells. **(E)** Max intensity projection of 30-minute timelapse videos of the basal layer of control, G1-stalled, and G2-enriched Ca^2+^-sensor mice (Rosa26-CAG-GCaMP6s; K14rtTA, Rosa26-CAG-GCaMP6s; K14rtTA; cdkn1b, and MMC-treated K14Cre; LSL-GCaMP6s). Time represented by color scale. Cdkn1b mice imaged 3 days post-induction. MMC mice imaged 2 days post-treatment. Scale bars: 25 μm. **(F)** Total Ca^2+^ events identified in G1-stalled cdkn1b mice, G2-enriched MMC-treated mice, and mitosis-enriched demecolcine-treated mice versus genetic or drug-treated controls. **** P *<*0.0001, Student’s t test, N = 8 control and 9 cdkn1b+ movies, ** P *<*0.01, One-way ANOVA, N = 8 DMSO control, 4 demecolcine, and 4 MMC movies. **(G)** PHATE time trajectory visualization of Ca^2+^ signaling in the cdkn1b+ G1-stalled basal layer and MMC-treated G2-enriched basal layer versus control basal layer shows disruption of smooth, directed and coordinated patterns of signaling in G1-stalled mice, whereas the G2-enriched basal layer maintains homeostatic patterns. **(H)** Representative persistence diagrams (*H*0: connected components, *H*1: loops, *H*2: voids) for G1- and G2-enriched conditions (R26-GCaMP6s; K14-rtTA; cdkn1b 3 days post-induction and MMC 2 days post-treatment). The persistence diagram from the G1-stalled group has more *H*1 (loop) features. **(I)** Wasserstein distances from the persistence diagrams of G2-enriched MMC, DMSO control, G1-stalled Cdkn1b, and genetic control mice. Distances are similar across G2-enriched and all control mice and different when compared to G1-stalled mice.

In order to better resolve the contributions of G2 versus M phase to Ca^2+^ signaling activity, we labeled all nuclei (K14-Cre; Rosa-CAG-LSL-GCaMP6s; K14-H2BmCherry) and tracked mitotic events as they occurred, while also recording Ca^2+^ signaling. Surprisingly, while extensive literature links transient elevations in Ca^2+^ with important steps of mitosis[49, 50, 51], we found that cells undergoing mitosis do not display cytosolic Ca^2+^ signaling during homeostasis (**Figure 3D, Movie 2**). To delve deeper into cells’ competence in propagating Ca^2+^ signals during mitosis, we treated mice with the drug demecolcine to enrich for mitotic basal cells and found that Ca^2+^ signals are abrogated (**Figure 3F, A4A, Movie 3**). Together, these data illustrated that G2 cells, but not mitotic cells, display spatiotemporally similar patterns of Ca^2+^ signaling, distinct from the very heterogeneous signaling patterns of G1 and S cells.

We next wondered whether G2 stem cells played any role in coordinating Ca^2+^ signaling and set out to deplete or enrich for G2 cells within the basal stem cell layer. First, we depleted G2 phases by stalling cells in the G1 state, using the Keratin-14 promoter to induce the cell cycle inhibitor Cdkn1b (p27) and combining this with a constitutive Ca^2+^-sensor (K14rtTA; pTRE-Cdkn1b; Rosa26-CAG-GCaMP6s). Through timelapse recordings, we observed a marked decrease in the amount of total Ca^2+^ signaling in these G1-stalled Cdkn1b mice compared to littermate controls (**Figure 3E, 3F, Movie 4**). Interestingly, we saw no change in the spatial patterns of Ca^2+^ signaling upon cell cycle inhibition (**Figure A4E**); however, our analyses revealed a temporal disruption, such that cells showed a trend towards a shorter maximal spike duration of their Ca^2+^ transients (**Figure A4F**). Comparison between G1-stalled and control cells at a population level via our GSTH analysis also revealed different patterns between the two groups (**Figure 3G, A6A**). The PHATE trajectories for the control group were smooth, consistent with wildtype tissue, suggesting the change of signals over the graph was generally steady; however, trajectories for the Cdkn1b positive group of G1-stalled cells showed different and more scattered patterns, with many loops and holes in the trajectories. These characteristics were also reflected in the persistence diagrams (based on the PHATE trajectories) and the corresponding Betti curves(**Figure 3H, A6C**). The G1-stalled datasets had more *H*1 features, representing loops that appear more often in rough trajectories. Betti curves for the G1-stalled group also showed that all loops are formed and closed at earlier thresholds, reflecting the scattered time trajectory and disruption in the spatiotemporal signaling coordination. These data demonstrate that in the absence of G2 cells, a homogeneous layer of G1-stalled cells is not able to carry out globally coordinated Ca^2+^ signaling.

Second, we tested the opposite scenario and enriched for G2 cells by treating with the drug Mitomycin C (MMC). Unlike mice enriched for G1 or mitotic cells, we observed normal local patterns of Ca^2+^ signaling in G2-enriched mice, similarly to DMSO vehicle-treated controls (**Figure 3E, 3F, Movie 5**). GSTH also revealed smooth PHATE trajectories (**Figure 3G, A6B**). As the final step of GSTH, to quantitatively compare the topology of the PHATE plots of G2-enriched signaling, we plotted persistence diagrams for each PHATE time trajectory (**Figure 3H**). The G2-enriched datasets showed *H*1 features (loops) that were formed and closed at later thresholds, reflecting the smoothness of the time trajectory and spatiotemporal coordination of the signaling, similar to controls. We next quantified Wasserstein distances between the persistence homology plots of MMC, Cdkn1b, and control populations (**Figure 3I**). Wasserstein or *earth mover’s distances* offer a powerful method to quantify differences between sets based on the cost of displacement from one configuration to another. A key advantage of the persistence diagram description of signaling is that Wasserstein distances are well-studied in this context[52]. The Wasserstein distances within the MMC G2-enriched and DMSO control groups were small, signifying similar PHATE trajectories (hence similar signaling patterns). However, the Wassertein distances between the Cdkn1b G1-stalled and genetic control groups were large, indicating different signaling patterns. Altogether, these data demonstrated that G1 cells require G2 cells to maintain Ca^2+^ signaling activity and coordinate signaling globally.

### Cx43 orchestrates Ca^2+^ signaling at large scales, but not across local neighborhoods, in the stem cell pool

So far, our work established a role for cell cycle phase in the regulation of Ca^2+^ signaling, yet we lack an understanding of the molecular mediators for Ca^2+^ propagation during epidermal regeneration. Gap junctions are known mediators of Ca^2+^ signaling in epithelial tissues, directly linking the cytoplasm of neighboring cells[53, 54]. To determine how cells within the basal stem cell layer are connected to their neighbors via gap junctions, we stained for Cx31 and Cx43, the two most highly expressed connexins in this layer[55, 56]. We observed high levels of Cx43 gap junctions compared to Cx31 (**Figure 4A**). Interestingly, we found that localization of Cx43 gap junctions to cellular junctions was highly heterogeneous from cell to cell, leading us to ask how it relates to the cell cycle states of this heterogeneous pool of stem cells. To this end, we examined Cx43 distribution via immunofluorescence at different stages of the cell cycle in the homeostatic epidermis of cell cycle reporter (Fucci2) mice. We found that stem cells display a gradient of junctional Cx43 expression as they pass through different cell cycle phases, with maximal junctional localization in the G1 stage (**Figure 4B, 4C, 4D**). These data suggest that Cx43 is dynamically regulated throughout the cell cycle and prompted us to interrogate Cx43 localization in the same genetic models and drug treatments from earlier. We found that enrichment or depletion of G1 cells confirms that Cx43 is most highly localized to cell-cell junctions in the G1 population (**Figure 4E**).

**Figure 4:**
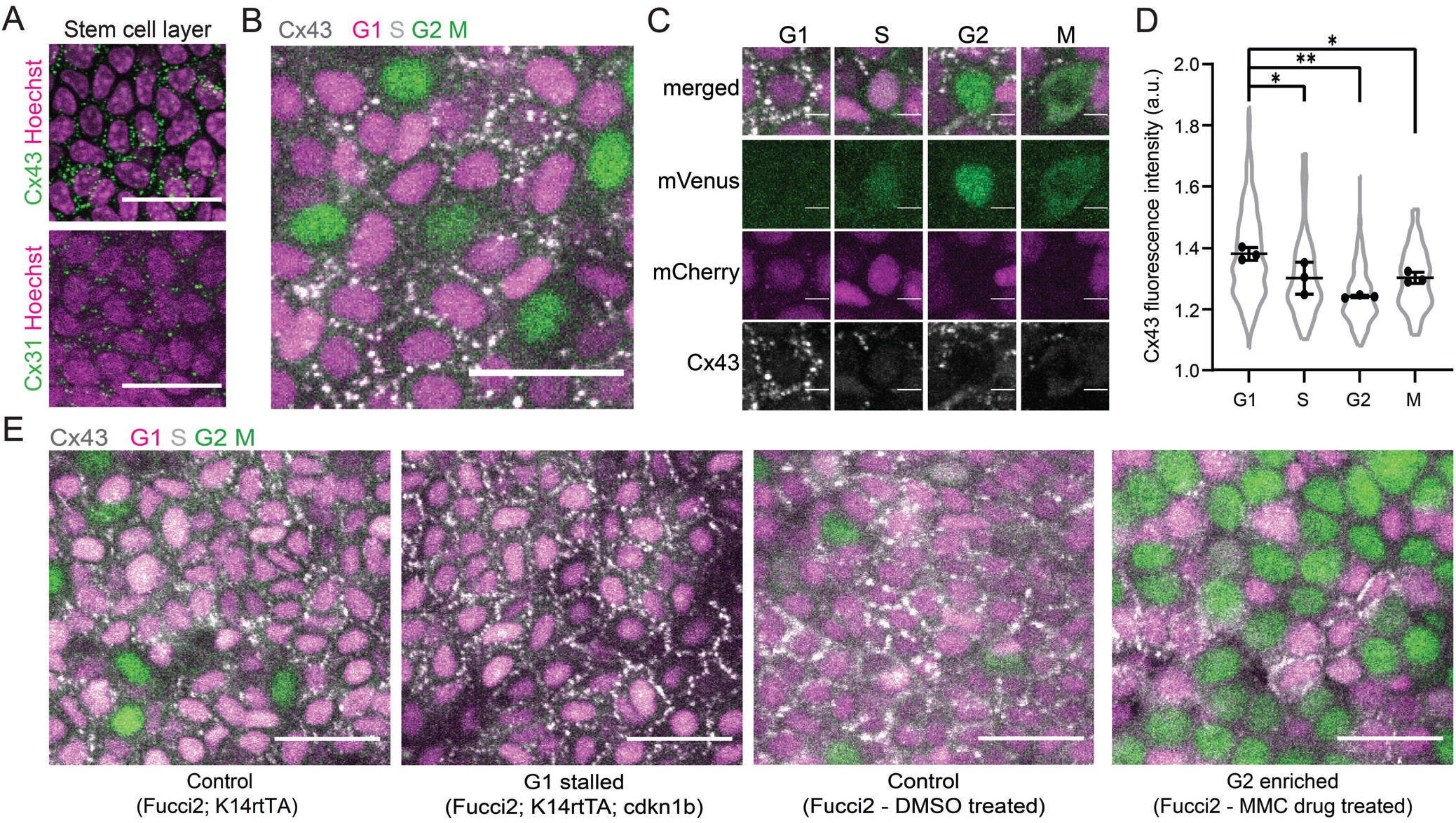
Cx43 distribution at junctions is enriched in G1 basal stem cells. **(A)** Maximum intensity projection of whole-mount immunoflu- orescence staining for Cx43 (top in green) and Cx31 (bottom in green) in the basal stem cell layer (Hoechst marking nuclei in magenta). Scale bars: 25 μm. **(B)** Maximum intensity projection of whole-mount immunofluorescence staining for Cx43, shown in white, in Rosa26p-Fucci2 mice, where G1 and S cells are mCherry^+^ (magenta) and S, G2, and M cells are mVenus^+^ (green). Scale bars: 25 μm. **(C)** Insets of G1, S, G2 and M cells in Rosa26p-Fucci2 mice with Cx43 immunofluorescence staining shown in white. Scale bars: 5 μm. **(D)** Quantification of Cx43 mean fluorescence intensity at the borders of G1, S, G2 and M cells in Rosa26p-Fucci2 mice. * P *<*0.05, ** P *<*0.01, One-way ANOVA, N = 3 control and 3 Cx43 cKO mice. **(E)** Cx43 whole-mount immunofluorescence staining (white) in G1- and G2-enriched Rosa26p-Fucci2 mice, where G1 and S cells are mCherry^+^ (magenta) and S, G2, and M cells are mVenus^+^ (green). G1-stalled and control (K14rtTA; cdkn1b; Rosa26p-Fucci2 and K14rtTA; Rosa26p-Fucci2) tissue was collected 3 days post-induction. G2-enriched MMC-treated and DMSO-treated control Fucci2 tissue was collected 2 days post-treatment. Scale bars: 25 μm.

Given G1 cells represent roughly 80% of the stem cell layer, we next asked whether Cx43 gap junctions are essential for homeostatic patterns of Ca^2+^ signaling within the heterogeneous basal stem cell layer. To address this, we crossed Cx43 conditional knockout mice (cKO) with a germline recombined Ca^2+^-sensor line (K14CreER; Cx43^fl/fl^; Rosa26-CAG-GCaMP6s and K14CreER; Cx43^+/+^; Rosa26-CAG-GCaMP6s littermate controls) and performed live timelapse imaging at 1-, 5-, and 7-days post-induction. We first confirmed loss of Cx43 protein expression within 5 days of recombination (**Figure A7A**). While loss of Cx43 abolishes Cx43 gap junctions, it does not completely abolish all gap junctions, as detected by immunofluorescence whole-mount staining for Connexin31 (**Figure A7B**).

Surprisingly, we observed no change in the average total number of Ca^2+^ events or the distribution of stem cell neighborhood sizes upon loss of Cx43 (**Figure 5A, 5B**). Temporal color coding frames of the timelapse movies from the Cx43 cKO mice revealed that spatially restricted neighborhoods of Ca^2+^ signaling oscillated more repeatedly within the 30-minute time period, in contrast to more dispersed and heterogeneous clustered signaling in the littermate controls (**Figure 5C, 5D, Movie 6**). Consequently, cells signaling in neighborhoods of 2 or more cells displayed an increased frequency of Ca^2+^ transients (**Figure 5E**). Loss of Cx43 resulted in a longer maximal spike duration for transients of Ca^2+^ signaling across all neighborhood sizes, most dramatically in single cells and small neighborhoods of 2 to 10 cells (**Figure 5F**). Disruption in the spatiotemporal characteristics of local neighborhoods of Ca^2+^ signaling demonstrated a modulatory role for Cx43 and prompted us to question whether loss of Cx43 affects signaling dynamics at a tissue-wide level.

**Figure 5:**
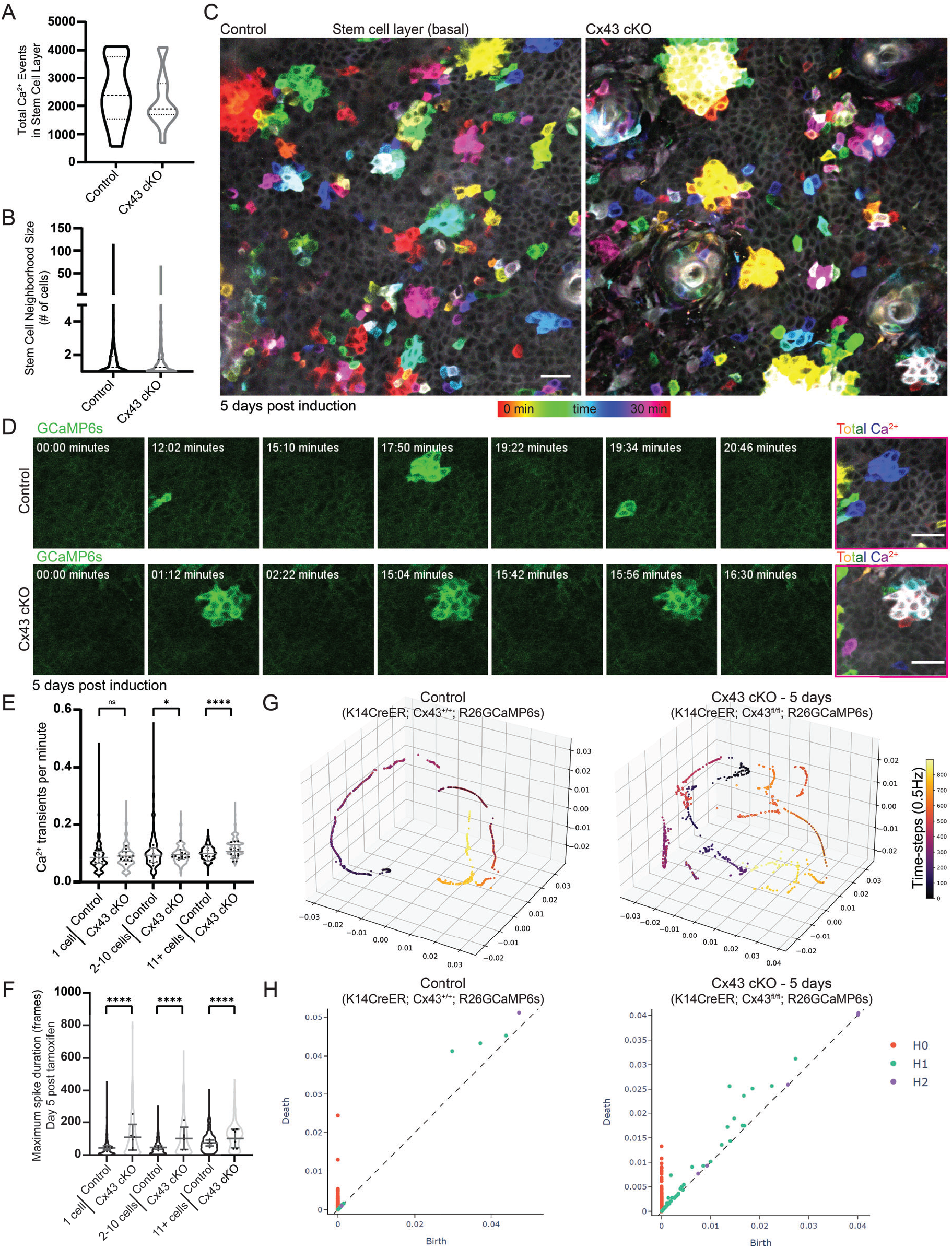
Cx43 is necessary for long-range Ca^2+^ signaling coordination but not local signaling activity. **(A)** Total number of Ca^2+^ signaling events in control versus Cx43 cKO mice. N = 11 (control) and 14 (Cx43 cKO) thirty-minute timelapse movies from at least 3 mice per condition. **(B)** Average neighborhood size of signaling in control versus Cx43 cKO mice. N = 11 (control) and 14 (Cx43 cKO) thirty-minute timelapse movies from at least 3 mice per condition. **(C)** Max intensity projection of 30-minute timelapse videos of the stem cell pool of control and Cx43 cKO Ca^2+^-sensor mice 5 days post-induction (Rosa26-CAG-GCaMP6s; K14-CreER; Cx43^+/+^ and Rosa26-CAG-GCaMP6s; K14-CreER; Cx43^fl/fl^). Color scale represents time. Repeated signaling manifests as white signal (the sum of colors). Scale bars: 25 μm. **(D)** Time-course of clustered signaling from 30-minute videos of the stem cell pool of control and Cx43 cKO Ca^2+^-sensor mice 5 days post-induction. Last image on right is max intensity projection with time represented by a color scale. Scale bars: 25 μm. **(E)** Ca^2+^ transients per minute per cell for three patterns of Ca^2+^ signaling (1 cell, 2-10 cells, or 11+ cells) in control versus Cx43 cKO Ca^2+^-sensor mice. NS for 1 cell, P = 0.0139 for 2-10 cells, P *<*0.0001 for 11+ cells comparison, Mann-Whitney test. **(F)** Maximal spike duration of Ca^2+^ transients per cell for three patterns of Ca^2+^ signaling (1 cell, 2-10 cells, or 11+ cells) in control versus Cx43 cKO mice 5 days post-tamoxifen induction. P *<*0.0001, Mann-Whitney test. **(G)** Representative PHATE visualization of Ca^2+^ signaling in the Cx43 cKO versus control stem cell pool shows disruption of smooth, directed and coordinated patterns of signaling in mice 5 days after loss of Cx43. **(H)** Representative persistence diagrams (*H*0: connected components, *H*1: loops, *H*2: voids) for control and Cx43 cKO mice (Rosa26-CAG-G1C2aMP6s; K14-CreER; Cx43^+/+^ and Rosa26-CAG-GCaMP6s; K14-CreER; Cx43^fl/fl^) 5 days post-induction. *H*1 features from Cx43 cKO mice appear later in time and have a longer persistence.

To address this, we applied GSTH to 30 minute timelapse movies from Cx43 cKO and control mice. We observed a striking loss of smooth, coordinated Ca^2+^ signaling time trajectories in the epidermal stem cell pool upon loss of Cx43 compared to littermate control mice (**Figure 5G, A8A**). Instead, Ca^2+^ signaling trajectories in the Cx43 mutant mice appeared more scattered and rough, showing more rapid changes of signals over the graph and less connected neighborhoods of intercellular signaling. This was also quantified and reflected by the persistence diagrams of each trajectory (**Figure 5H**). In these diagrams, *H*0 features represent connected components in the trajectory, *H*1 features represent loops, and *H*2 features represent voids. There were fewer *H*1 features from the persistence diagrams of the control group, and many of them were created and died at later stages. This indicates that signaling dynamics in the control group were less disjointed and formed smoother trajectories. Most *H*1 features in the Cx43 cKO group appeared and disappeared at earlier stages, thus these features were short-lived, reflecting rough trajectories. The differences in the persistence diagrams represent different topological features in the underlying PHATE trajectories, and therefore different Ca^2+^ signaling patterns after the loss of Cx43.

These differences were further revealed through topological descriptors such as Betti curves, which we depict for *H*1 features. The Betti curves for the control group show that all loops are formed and closed at later thresholds, while those in the Cx43 cKO exhibit loops that emerged at earlier stages (**Figure A8B**). These quantifications further demonstrate that loss of Cx43 leads to a disruption in the long-range signaling coordination we normally observe at homeostasis. Perturbed trajectories across the stem cell pool were evident as early as one day after loss of Cx43, suggesting a direct role for Cx43 in Ca^2+^ signaling regulation. As G1 cells make up the majority of the stem cell layer, Cx43 gap junctions allow for connectivity of these cells and coordination of long-range calcium signaling across the heterogeneous pool of stem cells in different cell cycle phases. Here we uncouple global from local communication and identify Cx43 as the molecular mediator necessary for signaling coordination across local signaling neighborhoods.

To finally understand the consequences of the loss of Cx43 and disruption of tissue-wide Ca^2+^ signaling coordination, we investigated the regenerative behaviors of cells in the stem cell pool. Basal stem cells are constantly balancing between two behaviors -self-renewal and differentiation. To determine whether loss of Cx43 affects these behaviors, we stained for the mitotic marker phospho-histone H3 (pH3) and for the early differentiation marker Keratin10 (K10)[57] in the Cx43 cKO mice 5 days post-induction. We observed a drop in bright pH3 staining (marking mitotic cells) in Cx43 cKO mice compared to littermate controls (**Figure A7C, A7D**), while punctate pH3 staining (small dots of staining across the nucleus, marking cells in the G2 stage of the cell cycle) was unchanged 5 days after loss of Cx43 (**Figure A7E**). This demonstrated that Cx43 is required for the progression of cells from G2 to mitosis. We did not see any change in the percent of K10 positive basal cells (**Figure A7F, A7G**), indicating that similar numbers of basal cells prepare to exit the basal layer on their differentiation trajectory despite a loss of Cx43. Additionally, we observed no noticeable change in basal cell density or overall epidermal thickness **(Figure A7H, A7I)** - two indicators of overall tissue homeostasis. These results suggest a possible downstream role for coordinated Ca^2+^ signaling via Cx43 gap junctions in regulating self-renewal within the stem cell pool.

## Discussion

As a general principle, regenerative tissues must orchestrate many types of cellular decisions across an ever-changing environment. Thus, communication among the tissue’s stem cells is paramount. Yet we still do not understand how the flow of information happens and at what scale. The necessity for Ca^2+^ signaling in many cellular processes has long been recognized. For example, the role for Ca^2+^ in proliferation, a fundamental property of regenerative tissues, was discovered more than 50 years ago[58]. However, the characteristics and regulation of Ca^2+^ signaling across a complex, multicellular regenerative tissue during homeostasis remains unclear due to significant challenges in studying both the spatiotemporal dynamics of the signaling, as well as the cellular complexity inherent to the tissue. In this study we set out to understand how stem cells communicate with one another via intercellular Ca^2+^ signaling and how they orchestrate this communication in the unperturbed stem cell pool. More broadly, we aim to understand how regenerative tissues integrate information flow across multiple scales to carry out essential homeostatic behaviors.

Our unique capacity to interrogate stem cell biology by resolving and tracking thousands of epidermal stem cells in live mice[59] in combination with newly developed unsupervised machine learning methods[32] enables fast live imaging and analysis of single cell Ca^2+^ dynamics in the adult mammalian skin epidermis. Development of these methods allowed us to overcome many of the challenges of gathering and processing higher dimensional signaling data across organisms and tissues. We observed a new unexpected paradigm of homeostatic Ca^2+^ signaling flow that is coordinated and nonrandom across the epidermal stem cell population, and identified two levels of regulation that coordinate local signaling patterns within the stem cell compartment and enable directed signal flow more globally. First, we observe that G2 cells are necessary for coordinated signaling activity, acting as signaling centers, while second, we find that neighboring G1 cells use Cx43 gap junctions to coordinate when and where intercellular Ca^2+^ signaling will occur across the whole stem cell compartment.

Most studies to date have used static analysis, perturbed conditions (such as skin explants), or *in vitro* models to study Ca^2+^ signaling within the epidermal stem cell pool[4, 60, 61, 62]. Our comprehensive quantification of local patterns of Ca^2+^ signaling within the basal stem cell compartment reinforced the existence of Ca^2+^ signaling activity within this pool of cells during homeostasis and without perturbation[63]. To understand whether Ca^2+^ signaling was spatially restricted to specific locations of the basal stem cell pool, we were able to revisit the same region of the tissue 24 hours later, observing that the whole stem cell pool is permissive to fluctuations in intracellular Ca^2+^ levels. These findings established that local patterns of Ca^2+^ signaling across neighborhoods of 1 to 10 cells are prevalent across a pool of heterogeneous stem cells, distinct from larger intercellular Ca^2+^ waves (ICWs) in epithelial sheets that are often a focus of attention in other systems.

Further, we were able to investigate the spread of information across multiple scales by developing the computational method, Geometric Scattering Trajectory Homology (GSTH)[39]. GSTH models the tissue as a cellular adjacency graph and derives descriptors for each timepoint using graph signal processing, allowing us to capture information about the underlying graph structure (in this case, the constituent stem cells) and how signals pass along graphs of different sizes and geometries, previously not possible to extract. These features are then aggregated into a trajectory using the dimensionality reduction method PHATE[40], allowing for understandable visualizations of complex data. While dimensionality reduction methods have been extensively and successfully applied to single cell -omics data to gain unique insights, they have yet to be widely applied to single cell signaling datasets (besides simple principle component analysis). GSTH then uses computational topology to quantify the whole trajectory and allow for comparisons between signaling dynamics of given variables, such as cell cycle stage. Importantly, GSTH, a data-driven, unsupervised approach, is highly versatile and can be applied in the future to imaging data with any molecular sensor and across tissues with widely different geometries.

GSTH revealed smooth trajectories along time for Ca^2+^ signaling data from the basal stem cell pool, revealing an emergent property of long-range spatiotemporally coordinated information flow. Because of the ability of GSTH to capture signaling patterns despite differences in tissue geometry, we were able to compare low-dimensional PHATE trajectories of Ca^2+^ signaling from the basal stem cell pool and the neuronal visual cortex. Both stimulated and spontaneous activity within the neuronal visual cortex displayed chaotic Ca^2+^ signaling trajectories, highlighting the unique cohesiveness of the epithelial basal layer. These applications of GSTH to explore and understand the dynamic processes of stem cells open a new realm of possibilities for understanding not only the molecular signaling pathways that underpin regenerative processes in homeostatic and perturbed tissue environments, but also any cellular signaling system beyond regenerative ones.

While Ca^2+^ signaling has been linked to cell proliferation in developmental and regenerative contexts[64, 65], the relationship between cell cycle progression and Ca^2+^ signaling in living animals is poorly understood. We found that when cells are in G2, they display Ca^2+^ signaling patterns that are more similar to each other across spatial and temporal dimensions than when they are in G1 or S. Additionally, these G2 cells are essential to tissue-level signaling coordination, in the sense that when cells are depleted of G2 cells, smooth signaling dynamics are disrupted and overall signaling activity is lower. Finally, in contrast to the historical paradigm that implicates Ca^2+^ signaling during mitosis in cultured cells and early embryos[49, 50, 51], we find that mitotic cells do not participate in Ca^2+^ signaling. This is consistent with more recent reports describing cells’ inability to activate store-operated calcium entry during mitosis[46, 47].

The dynamic regulation of Cx43 localization at cell-cell junctions as cell progress through the cell cycle reveals a new mechanism for coordinating the flow of Ca^2+^ across thousands of cells. There is some limited evidence showing the cell cycle-dependent expression of other Ca^2+^ signaling pathway proteins, such as Orai2 and voltage-gated Ca^2+^ channels[45, 66]. This finding opens the question of whether other signaling proteins could be dynamically regulated throughout the cell cycle and how cells are able to carry out this regulation.

Our discovery that Cx43, a molecular player implicated in the past as an on/off switch for intercellular Ca^2+^ signaling, is necessary for long-range coordination of Ca^2+^ signaling across the stem cell compartment but not for local clusters of Ca^2+^ transients revealed a complex role for Cx43 gap junctions in this compartment. This is in contrast to other work in this context, which has often used drug treatments to target gap junctions and then looked at Ca^2+^ signaling dynamics, showing a global disruption of intercellular Ca^2+^ signaling. Therefore, our results open a new way of thinking about this and suggest that Cx43 plays a unique role in coordinating information flow across this community of stem cells. Other molecular regulators, including other gap junction proteins such as Cx31, may compensate for the loss of Cx43 to carry out local patterns of Ca^2+^ signaling.

After the loss of Cx43, we saw that basal cells expressed early differentiation markers at a normal rate. However, stem cells across the basal layer were not able to properly compensate for neighbor loss at mitotic rates observed during homeostasis. Because we are using a model with fine temporal control, we capture the more immediate effects of loss of Cx43 on tissue homeostasis. In other instances, constitutive loss or alteration of wildtype Cx43 expression, including in the context of the disease Oculodentodigital dysplasia (ODDD), does not cause apparent phenotypes in the skin[67, 68]. This could mean the epidermal stem cell pool is eventually able to compensate for loss of Cx43 over time. Others have recently proposed a similar role for gap junction-mediated Ca^2+^ signaling in regulating the balance of regenerative behaviors in the *Drosophila* blood progenitor and intestinal stem cell pools[31, 65]. This might suggest that Cx43 gap junctions allow Ca^2+^ signaling to act as a common signaling mechanism across tissues to balance regenerative behaviors.

To our knowledge, our study represents the first time that stem cell progression through cell cycle and Ca^2+^ signaling have been studied in conjunction with one another *in vivo*. Additionally, while analysis of individual epidermal stem cells gives the impression of localized and random bursts across neighborhoods of 1 to 10 cells, our global analysis shows an underlying long-range and time-directed coordination of Ca^2+^ signaling across thousands of cells. Together, our results provide insight into how a heterogeneous pool of stem cells have different roles, coming together as a community to regulate a molecular pathway at large scale to maintain proper homeostasis, new concepts to the fields of both regenerative biology as well as calcium and signaling biology fields. Further, we have shown that our GSTH pipeline can be widely used to further interrogate stem cells with any kind of spatially and temporally patterned signaling dynamics — opening up this type of study to other signaling paradigms and tissues beyond regenerative ones.

## Experimental Methods

### Mice and experimental conditions

K14-Cre[69] mice were obtained from E. Fuchs (Rockefeller University). R26p-Fucci2[48] mice were obtained from S. Aizawa (RIKEN). K14-H2BmCherry mice were generated in the laboratory and described previously[70]. Cx43^fl/fl^[71, 72], Rosa26-CAG-LSL-GCaMP6s[73], mTmG[74], Sox2-Cre[75], K14-CreER[69], K14-rtTA[76], and tetO-Cdkn1b[77] mice were obtained from The Jackson Laboratory. Germline recombined Rosa26-CAG-GCaMP6s mice were generated by crossing Rosa26-CAG-LSL-GCaMP6s to Sox2-Cre mice. To block the cell cycle progression of epithelial cells during G1, Rosa26-CAG-GCaMP6s mice were mated with K14-rtTA; tetO-Cdkn1b mice (Rosa26-CAG-GCaMP6s; K14-rtTA; tetO-Cdkn1b) and given doxycycline (1 *mg*·*ml*^−1^) in potable water with 1% sucrose between P45 and P60. Doxycycline treatment was sustained until imaging was performed three days later. Siblings without the tetO-Cdkn1b allele (Rosa26-CAG-GCaMP6s; K14-rtTA) were used as controls. Mice from experimental and control groups were randomly selected from either sex for live imaging experiments. No blinding was done. All procedures involving animal subjects were performed under the approval of the Institutional Animal Care and Use Committee (IACUC) of the Yale School of Medicine.

### *In vivo* imaging

Imaging procedures were adapted from those previously described[26, 27]. All imaging was performed in distal regions of the ear skin during prolonged telogen, with hair removed using depilatory cream (Nair) at least 2 days before the start of each experiment. Mice were anaesthetized using an isoflurane chamber and then transferred to the imaging stage and maintained on anesthesia throughout the course of the experiment with vaporized isoflurane delivered by a nose cone (1.25% in oxygen and air). Mice were placed on a warming pad during imaging. The ear was mounted on a custom-made stage and a glass coverslip was placed directly against it. Image stacks were acquired with a LaVision TriM Scope II (LaVision Biotec) laser scanning microscope equipped with a tunable Two-photon Vision II Ti:Sapphire (Coherent) Ti:Sapphire laser and tunable Two-photon Chameleon Discovery Ti:Sapphire laser (Coherent) and Imspector Pro (LaVision Biotec, v.7.0.129.0). To acquire serial optical sections, a laser beam (940*nm*, 1120*nm* for mice and whole-mount staining) was focused through a 20x or 40x water-immersion lens (NA 1.0 and 1.1 respectively; Zeiss) and scanned with a field of view of 500 *μm*^2^ or 304 *μm*^2^, respectively at 800 Hz or through a 25x water-immersion lens (NA 1.0; Nikon) and scanned with a field of view of 486 *μm*^2^ at 800 Hz. Z-stacks were acquired in 0.5–3 μm steps to image a total depth of up to 100 μm of tissue. To visualize large areas, 2–64 tiles of optical fields were imaged using a motorized stage to automatically acquire sequential fields of view. Visualization of collagen was achieved via the second harmonic signal at 940*nm*. For all time-lapse movies, the live mouse remained anesthetized for the length of the experiment and serial optical sections were captured at intervals of 2 seconds. For revisits, the same region of live mouse skin was imaged across intervals of multiple days. Anatomical features and patterns of hair follicles and collagen were used as landmarks for finding the same skin location (see **Figure A1A**).

### Image Analysis

Raw image stacks were imported into FIJI (ImageJ, NIH) for analysis. Individual optical planes, summed or max Z stacks of sequential optical sections were used to assemble figures. To prepare movies where the nuclear signal bleached over the course of the timelapse, we used the Fiji Bleach Correction plugin[78], specifying the Simple Ratio Method.

Segmentation of actively signaling cells was performed using the CaImAn MATLAB package as previously described[33]. In order to segment all cells in the field of view, including non-flashing cells, we used part of the MATLAB package from Romano et al[29], a watershed segmentation method. We normalized the fluorescence intensity of each cell at each timepoint to the minimum fluorescence intensity of that cell as a baseline. From the normalized fluorescence values for each segmented cell, we used peak finding in MATLAB (version R2018b) and then fit Gaussian curves to each peak to be able to quantify spike duration, peak intensity, frequency of flashing, etc. To quantify the neighborhood size of clustered signaling, we created a graph for each timelapse, where each node represented one segmented, spiking cell. We connected nodes that represented cells spiking directly adjacent to one another (spatial neighbors) within 10 seconds of each other (temporally correlated). We then counted the number of connected nodes to quantify the size of each signaling neighborhood.

### Whole-mount staining

Ear tissue was incubated epidermis side up in 5 *mg*·*ml*^−1^ Dispase II solution (Sigma, 4942078001) at 37 °C for 15 min, and epidermis was separated from dermis using forceps. The epidermis was fixed in 4% paraformaldehyde in PBS for 15 min at room temperature, washed and blocked with 0.2% Triton X-100, 5% normal donkey serum, 1% BSA in PBS. The samples were then incubated with primary antibodies for 12 h at 4 degrees and with secondary antibodies for approximately 2 hours at room temperature. Primary antibodies used were as follows: purified mouse anti-Connexin 43, C-terminal, clone P4G9 (1:100, Sigma, MABT901), rabbit anti-Connexin 30.3 polyclonal antibody (1:100, ThermoFisher, 40-0900), rabbit anti-Connexin 26 polyclonal antibody (1:100, ThermoFisher, 71-0500), rabbit anti-Connexin 31 polyclonal antibody (1:100, ThermoFisher, 36-5100), guinea pig anti-K10 (1:200; Progen, GP-K10), rabbit anti-pH3 (1:300; Millipore, 06-570). All secondary antibodies used were raised in a donkey host and were conjugated to AlexaFluor 488, 568, or 647 (Thermofisher). Some tissue was then incubated with Hoechst 33342 (Becton Dickinson; H3570, 1:500) for 15 min, then washed with blocking solution. Finally, the tissue was mounted with Vectashield Anti-fade mounting medium (Vector Laboratories) or SlowFade™ Diamond Antifade Mountant (ThermoFisher) and a #1.5 coverslip and imaged on a LaVision TriM Scope II as described in ‘In vivo imaging’.

### Tamoxifen Induction

To induce expression of membrane-GFP and/or loss of Cx43 expression, K14-CreER; Cx43^fl/fl^; mTmG mice or K14Cre-ER; Cx43^fl/fl^ mice were given three doses of Tamoxifen (20*mg/kg* body weight in corn oil) 3, 4, and 5 days before imaging or tissue collection by intraperitoneal injection. In order to observe phenotypes of total loss of Cx43 just one day after recombination, we also topically applied 0.01 mg (Z)-4-Hydroxytamoxifen (4-OHT) in an ethanol-Vaseline slurry to the ear of Rosa26-CAG-GCaMP6s; K14CreER; Cx43^fl/fl^ or Rosa26-CAG-GCaMP6s; K14CreER; Cx43^+/+^ mice one day before the start of imaging.

### Topical drug treatment

To stall cells as they transition from S to G2 phase of their cell cycles, Mitomycin C (MMC)[79] was delivered topically to the ear skin. MMC was dissolved in a 15 *mg*·*ml*^−1^ stock solution in dimethyl sulfoxide (DMSO) and then diluted 100 times in 100% petroleum jelly (Vaseline; final concentration is 150 *mg*·*ml*^−1^). One hundred micrograms of the mixture of the MMC and the petroleum jelly was spread evenly on the ear 1 and 2 days before imaging. A mixture of 100% DMSO in petroleum jelly (1:100) was used as a vehicle control. Demecolcine was used to block microtubule polymerization[80]. Colcemid was dissolved to 25 *mg*·*ml*^−1^ stock solution in DMSO and delivered as described for the MMC treatment.

### Statistics and reproducibility

Biostatistical analyses were performed using GraphPad Prism (version 9.2) software (GraphPad Inc., La Jolla, CA). Statistical comparisons were made using an unpaired two-tailed Student’s t test, Mann-Whitney test, or the one-way analysis of variance (ANOVA) with multiple comparison’s test. Differences between the groups were considered significant at P *<* 0.05, and the data are presented as means ± standard deviation unless otherwise noted.

### Computational Methods

#### Diffusion Geometry

A useful assumption in representation learning is that high dimensional data originates from an intrinsic low dimensional manifold that is mapped via nonlinear functions to observable high dimensional measurements; this is commonly referred to as the manifold assumption. Formally, let ℳ^*d*^ be a hidden *d*-dimensional manifold that is only observable via a collection of *n* ≫ *d* nonlinear functions *f*_1_,…, *f*_*n*_ : ℳ^*d*^ → ℝ that enable its immersion in a high dimensional ambient space as *F* (ℳ^*d*^) = *{***f** (*z*) = (*f*_1_(*z*),…, *f*_*n*_(*z*))^*T*^ : *z* ∈ ℳ^*d*^} ⊆ ℝ^*n*^ from which data is collected. Conversely, given a dataset *X* = {*x*_1_,…, *x*_*N*_}⊂ ℝ^*n*^ of high dimensional observations, manifold learning methods assume data points originate from a sampling 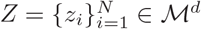 of the underlying manifold via *x*_*i*_ = **f** (*z*_*i*_), *i* = 1,…, *n*, and aim to learn a low dimensional intrinsic representation that approximates the manifold geometry of ℳ^*d*^.

To learn a manifold geometry from collected data, scientists often use the diffusion maps construction of [38] that uses diffusion coordinates to provide a natural global coordinate system derived from eigenfunctions of the heat kernel, or equivalently the Laplace-Beltrami operator, over manifold geometries. This construction starts by considering local similarities defined via a kernel 𝒦 (*x, y*), *x, y* ∈*F* (ℳ^*d*^), that captures local neighborhoods in the data. We note that a popular choice for 𝒦 is the Gaussian kernel exp(−‖ *x* − *y*‖ ^2^/σ), where σ *>* 0 is interpreted as a user-configurable neighborhood size. However, such neighborhoods encode sampling density information together with local geometric information. To construct a diffusion geometry that is robust to sampling density variations, we use an anisotropic kernel

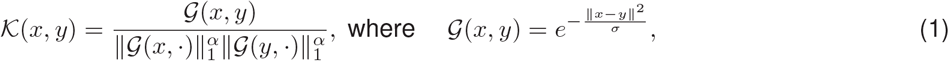

as proposed in [38], where 0 ≤ *α* ≤1 controls the separation of geometry from density, with *α* = 0 yielding the classic Gaussian kernel, and *α* = 1 completely removing density and providing a geometric equivalent to uniform sampling of the underlying manifold. Next, the similarities encoded by 𝒦 are normalized to define transition probabilities 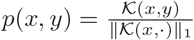 that are organized in an *N* × *N* row stochastic matrix

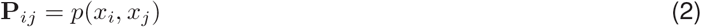

that describes a Markovian diffusion process over the intrinsic geometry of the data. Finally, a diffusion map [38] is defined by taking the eigenvalues 1 = *λ*_1_ ≥ *λ*_2_ ≥ · · · ≥ *λ*_*N*_ and (corresponding) eigenvectors 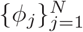 of **P**, and mapping each data point *x*_*i*_ ∈ *X* to an *N* dimensional vector 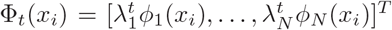, where *t* represents a diffusion-time (i.e., number of transitions considered in the diffusion process). In general, as *t* increases, most of the eigenvalues 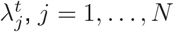, become negligible, and thus truncated diffusion map coordinates can be used for dimensionality reduction [38].

Note that thus far we have described the diffusion operator construction in the case when datapoints are sampled from a high dimensional space. However, in some cases the connectivity structure of the datapoints may be more apparent then their ambient dimensions. This is true for cases where the data already *comes as a graph* in the case of social networks or protein interactions. In our case too, the data can be easily turned into a connectivity structure on the basis of spatial adjacency. In these cases, the distance computation is not necessary and one can simply start with the adjacency or connectivity structure.

#### Cellular Graphs and Graph Signals

We represent the imaged tissue as a graph *G* ={*V, E*}, consisting of nodes *v*_*i*_ ∈ *V* and edges (*v*_*j*_, *v*_*k*_) ∈ *E*, where each node *v*_*i*_ represents a cell and a pair of nodes *v*_*j*_ and *v*_*k*_ is connected with an edge based on a predefined criterion. For epithelial cells, we connect nodes that are spatially adjacent (within 2 μm of each other), as the flow of signals is thought to be between spatially proximal cells. On the other hand, neurons can have long processes that are often hard to image, and therefore we use correlation between neurons’ Ca^2+^ signals to connect the neuronal graph. Finally, the connectivity of graph *G* can be described by its adjacency matrix **A**, where **A**_*ij*_ = 1 if *v*_*i*_ and *v*_*j*_ are connected and 0 otherwise. The degree of each vertex is defined as a diagonal matrix **D**, where 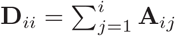.

Graph signals can associate with each node or edge in a graph. In the Ca^2+^ signaling data, the signals associated with cell *v*_*i*_ is the normalized Ca^2+^ fluorescence intensity at each timestep *t*. Since every cell has a related Ca^2+^ signal, this signal *X*(*v*_*i*_, *t*) is defined over the whole graph for timestep *t*.

#### Geometric scattering for timepoint embeddings

The geometric scattering transform is an unsupervised method for generating embeddings for graph-structured data[39]. It is constructed by applying a cascade of graph wavelet transforms followed by a nonlinear modulus operation such as an absolute value nonlinearity[39, 81]. Graph wavelets are designed based on the diffusion operator (lazy random walks) 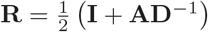 over a graph, i.e.,

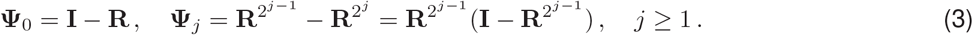

The multi-scale nature of graph wavelets allows the geometric scattering transform to traverse the entire graph in one layer, which provides both local and global graph features. Summation of the signal responses is used to obtain invariant graph-level features. Since the summation operation could suppress the high frequency information, it could be complemented by using higher order summary statistics of signal *x*. Due to the iteration of applying graph wavelets followed by a nonlinaer modulus operation, as shown in **Figure 2**, geometric scattering transforms can be constructed as in a multi layer (or multi order) architecture. Specifically, the zeroth-order scattering coefficients are calculated by taking statistical moments of the summation of signals, and the first order features are obtained by applying a graph wavelet, which aggregates multiscale information of the graph. Second-order geometric scattering features can further augment first order features by iterating the graph wavelet and absolute value transforms. The collection of graph scattering features provides a rich set of multiscale invariants of the graph G and can be used under both supervised and unsupervised settings for graph embedding.

For a signal *X*(*t*_*i*_) = [*X*(*v*_1_, *t*_*i*_),*X*(*v*_2_, *t*_*i*_),…,*X*(*v*_*m*_, *t*_*i*_)] we compute the zeroth-order scattering coefficients for each vertex/cell for timepoint *t*_*i*_ as follows:

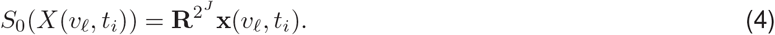

The diffusion operator (lazy random walks) **R** here works as a low pass filter as shown in [39] and provides local averaging of neighboring cell patterns[82]. Unlike the summation operator that averages all vertex information and suppresses the high frequency information and hence has to be retrieved by higher order statistical moments, this retains finer description of cell/vertex embeddings. Then, by concatenating the wavelet coefficients for each cell/vertex at timepoint *t*_*i*_, we can obtain the corresponding timepoint embedding *S*_0_(*X*(*t*_*i*_)) for timepoint *t*_*i*_. Finally, the timepoint embedding for *N* timepoints can be calculated and the resulting *S*_0_(*X*(*t*)) = {*S*_0_(*X*(*t*_0_)), *S*_0_(*X*(*t*_1_)),…, *S*_0_(*X*(*t*_*n*_))} is a feature matrix of dimension *N* × *M* , where *N* is the number of timepoints and *M* is the number of cells. We hence obtain the zeroth-order scattering coefficients for the *N* timepoints. The scattering transform here is a result of local averaging of wavelet coefficients.

As in [39], the zeroth-order scattering features can be augmented by first-order scattering features by applying graph wavelets and extracting finer description of high frequency response of a signal *X*(*t*_*i*_). Specifically, the first-order scattering coefficients for each time point at each vertex/cell are calculated as

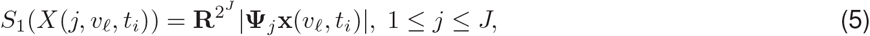

The value **Ψ**_*j*_**x**(*v*_𝓁_, *t*_*i*_) aggregates the signal information **x**(*v*_*m*_, *t*_*i*_) from the vertices *v*_*m*_ that are within 2^*j*^ steps of *v*_𝓁_ . It responds to sharp transitions or oscillations of the signal **x** within the neighborhood of *v*_𝓁_ with radius 2^*j*^ (in terms of the graph path distance). By concatenating all the vertex/cell embeddings, we can obtain the first order scattering coefficients *S*_1_(*X*(*t*_*i*_)) for timepoint *t*_*i*_.

Finally, the second-order scattering coefficients can be obtained by further applying graph wavelets and extract even finer description of high frequency response of the signal *X*(*t*_*i*_):

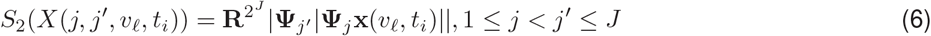

The above calculations are conducted for each timepoint and a total of *N* timepoints. The first-order and second-order scattering transform will generate a feature matrix of shape *N* ×(*M* × *J*) and 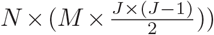, respectively, as timepoint embeddings for the *N* timepoints. Finally, the zeroth-order, first-order and second-order scattering coefficients were combined together as the embeddings for each time point *S*(*X*(*t*_*i*_)). The scale of the wavelet *J* was selected based on the diameter of graphs, and the number of scattering coefficients generated depended on the graph sizes.

## PHATE

PHATE is a dimensionality reduction method that captures both local and global nonlinear structure through constructing a diffusion geometry[40]. It computes the diffusion operator as in Equation 2. However, rather than eigendecomposing this operator to find new coordinates, PHATE creates a new distance matrix from **P** by defining an M-divergence between datapoints, called *potential distance* as 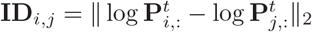 between corresponding *t*-step diffusion probability distributions of the two points.

The advantage of this step is that the information theoretic distance between probabilities emphasizes differences in lower probabilities (corresponding to distant points) as well as high probabilities (corresponding to neighbors), and therefore globally contextualizes the point. The resulting information distance matrix **ID** is finally embedded into a low dimensional (2D or 3D) space by metric multidimensional scaling (MDS), and makes it possible to visualize intrinsic geometric information from data. In [40], authors demonstrate that PHATE performs better than all compared methods including diffusion maps and UMAP in preserving *denoised manifold affinity (DeMAP)* in low dimensions and, in particular, excels at preserving trajectory structures without shattering.

### PHATE trajectories of timepoint embeddings

The time point embeddings *S*(*X*(*t*_*i*_)) from geometric scattering form a matrix of dimensions *T* × *M* , where *T* is the number of time points in the data and *M* is the number of scattering coefficients for each time point. We can visualize these embeddings by applying PHATE. Following our previous description of PHATE, we calculated a distance matrix **D** = ‖*S*(*X*(*t*_*i*_)) − *S*(*X*(*t*_*j*_)‖_2_ based on the Euclidean distance between time point embeddings and applied an *α*-decaying kernel *K* with a locally-adaptive bandwidth *ϵ*_*k,i*_ corresponding to the *k*-NN distance of the *i*-th data point to generate an affinity matrix **W** as well as the diffusion operator **P**. The elements of **W** are given by:

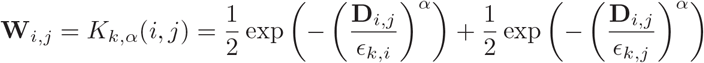

The decaying factor *α* regulates the decay rate of the kernel (smaller *α* ⇒ kernel with lighter tails), *α* = 2 corresponding to the Gaussian. The diffusion operator **P** can then be obtained by calculating the row-sum of the affinity matrix **W** with element **P**_*i,j*_ giving the probability of moving from the *i*-th to the *j*-th data point in one time step. The global structure of the data can be further learned through calculating the *t*th power of the diffusion operator **P**, which propagates affinity of the data through diffusion up to a scale of *t*. The optimal value *t* for diffusion is automatically chosen to be the knee point of the von Neumann entropy of **P**^*t*^. This diffusion operator is then log scale transformed and converted to a potential distance matrix *ID*((*X*)) which is embedded by MDS to result in 3-D PHATE embedding coordinates *E*(*t*) = (*E*_1_(*X*(*t*)), *E*_2_(*X*(*t*)), *E*_3_(*X*(*t*))) for each time point *t*, and point cloud *E* = {*E*(*t*_1_), *E*(*t*_2_),… *E*(*t*_*n*_)}.

The 3D coordinates enable visualization of the trajectory, which reflects the time-varying patterns of Ca^2+^ fluores-cence data. Thus neighbors in the PHATE embedded trajectories indicate similar signaling patterns even if they occur at distal timepoints. In fact, many of the dynamics we notice have circularity, which motivates the use of topology in the next section.

### Persistent homology and topological data analysis

Topological data analysis (TDA) refers to techniques for understanding complex datasets by their topological features, i.e., their connectivity[83]. Here we focus on the topological features of a data graph where the simplest set of topological features are given by the number of connected components *b*_0_ and the number of cycles *b*_1_, respectively. Such counts, also known as the Betti numbers, are coarse graph descriptors that are invariant under graph isomorphisms. Their expressivity is increased by considering a function *f* : *V* × *V* →*ℝ* on the vertices of a graph *G* = (*V, E*) with vertex set *V* and edge set *E*. Since *V* has finite cardinality, so does the image im *f* , i.e., im *f* = {*w*_1_, *w*_2_,…, *w*_*n*_} . Without loss of generality, we assume that *w*_1_ ≤…≤*w*_*n*_. We write *G*_*i*_ for the subgraph induced by filtering according to *w*_*i*_, such that the edges satisfy 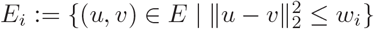. The subgraphs *G*_*i*_ satisfy a nesting property, as *G*_1_ ⊆ *G*_2_ ⊆ … ⊆ *G*_*n*_. When analyzing a point cloud, the vertices of each *G*_*i*_ arise from spatial coordinates for the data and *w*_*i*_ constitutes a distance threshold between points, such that *G*_*n*_ is a fully-connected graph, containing all the vertices from *V* . This is commonly known as the Vietoris-Rips (VR) filtration.

It is then possible to calculate topological features alongside this *filtration* of graphs, tracking their appearance and disappearance as the graph grows. If a topological feature is created in *G*_*i*_, but destroyed in *G*_*j*_ (it might be destroyed because two connected components merge, for instance), we represent this by storing the point (*w*_*i*_, *w*_*j*_) in the *persistence diagram* 𝒟_*f*_ associated to *G*. Another simple descriptor is given by the Betti curve of dimension *d* of a diagram 𝒟, which refers to the sequence of Betti numbers of dimension *d* in 𝒟, evaluated for each threshold *w*_*i*_.

#### Persistent homology analysis of PHATE trajectories

In this study, to obtain an invariant characterization of the generated PHATE trajectories *E* for topological data analysis, we calculated their persistent homology. Specifically, we calculated the persistent homology of *E* via a Vietoris–Rips filtration *V R*_*s*_(*E*). The Vietoris–Rips complex of *E* is defined as the filtered simplicial complex that contains a subset of *E* as a simplex if and only if all pairwise distances in the subset are less than or equal to *s*, i.e., *V R*_*s*_(*E*) ={{*n*_0_, …, *n*_*m*_}| ∀*i, j d*(*i, j*) ≤ *s*}. We noted here that we could also use the potential distance *ID* from PHATE, however we directly used the PHATE coordinates and the Euclidean distance for simplicity.

As described above, from *V R*_*s*_(*E*), we obtain a set of persistence diagrams *Q* consisting of birth-death-dimension triples [*b, d, q*] that describe multiscale topological features of *E*. Each such point corresponds to a topological feature in the trajectory, which appears at a certain birth time and disappears at a death time. Note that the times are supposed to be understood with respect to the parameter *s* from above. A point’s distance from the diagonal therefore represents the prominence or the eponymous *persistence* of the associated topological feature; higher values indicate that the feature occurs over a large scale, thus increasing its significance. We further calculated the associated Betti curves for each *Q*, resulting in a simple summary curve *B*(*Q, q*) for the *q*th dimension consisting of the number of points (*b*_*i*_, *d*_*i*_) in *Q* such that *b*_*i*_ ≤ *s* < *d*_*i*_. The Betti curve characterizes the connectivity of *V R*_*s*_(*E*) and, by extension, of the Ca^2+^ fluorescence data.

### Synthetic dataset for timepoint embeddings

To validate the utility of our method, we first tested it on three synthetic datasets we created, which simulated different signal diffusing scenarios.

We took a graph *G* created from one of our Ca^2+^ signaling samples with 1867 vertices (cells) and used a normalized graph Laplacian *L* to diffuse a Dirac signal *x* defined on node *i*, where *x*_*i*_ = 1 and 0 elsewhere. We diffused the signal over the graph for 300 steps via:

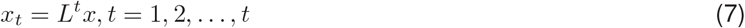

This resulted in a series of signals *X* = {*x*_1_, *x*_2_,…, *x*_*t*_} of 300 timesteps, each more diffused than the previous.

#### Synthetic testcase 1

We first added normalized random noise *ϵ* ∼ 𝒩 (*μ*, σ^2^) with *μ* = 0 and σ = 0.001 to *X* and obtained perturbed signals *X*_*perturbed*_, with *X*_*perturbed*_ = *X* + *ϵ*. The individual instances of *X*_*perturbed*_ are thus similar but not exactly the same as the original signals. We then combined *X* and *X*_*perturbed*_ to form a new 600-step series of signals.

#### Synthetic testcase 2

Next we created another signal *x*′ similar to the previously defined Dirac signal *x* centered on node *i*. This new signal *x*′ is centered on both node *i* and node *j*. In other words, 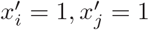 and 0 otherwise. Therefore, the diffusion of this new signal *x*′ on the graph initially was similar to signal *x*, but eventually diffused to different patterns. We also diffused this signal *x*′ for 300 steps and obtained another signal *X*′. As in the previous testcase, we combined *X* and *X*′ to form another series of signals of 600 steps.

#### Synthetic testcase 3

Finally, we took the first 50 timesteps from *X*, then starting from timestep 51, we created new signals for each timestep. Specifically, we first removed all signals defined on each cell, then 100 cells were randomly picked to choose one of three signals 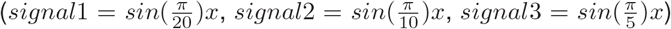 to spike for a random interval of 11 timesteps during a total of 550 timesteps. These signals were only defined on each cell and not diffused to other cells. Finally, we combined the 50 timesteps from *X* with the newly generated signals and formed a series of 600 timesteps.

### Comparison of GSTH with PHATE, t-SNE, PCA and UMAP

We compare application of the proposed GSTH method on the three synthetic datasets with approaches that ablate or replace steps in the GSTH method. In particular, we test:

- Applying PHATE directly on the raw input signals to obtain time-trajectories—without the use of the geometric scattering transform.
- Applying PCA on the generated scattering coefficients instead of PHATE.
- Applying t-SNE on the generated scattering coefficients instead of PHATE.
- Applying UMAP on the generated scattering coefficients instead of PHATE.

For the synthetic testcase 1, we aim to compare the approaches for their stability to small perturbations as well as their ability to retrieve signal diffusion dynamics on the graph. As shown in **Figure A9A**, after applying GSTH, time points with perturbed signals overlapped with time points with original signals, showing scattering transform and PHATE are invariant to small degrees of noise. The smooth trajectory also reflects that the scattering transform and PHATE of GSTH can effectively capture the signal propagation on the graph. By contrast, directly using PHATE on the raw input signals will result in the condensed timepoints in **Figure A9C**, thus failing to retrieve the dynamics. While applying PCA (**Figure A9D**) and t-SNE (**Figure A9E**) on the generated scattering coefficients can retrieve the dynamics to some extent, **Figure A9D** shows a more disrupted trajectory and the trajectory from **Figure A9E** overlaps with itself. Similarly, applying UMAP (**Figure A9F**) on the generated scattering coefficients also led to overlapping timepoints. All these methods thus failed to reflect the propagation of signals on the graph.

For the second synthetic dataset, we further compare the ability of different approaches to retrieve signal diffusion dynamics on the graph under a more complex scenario. For GSTH (**Figure A9H**) time points from two signal sources formed two branches with their starting points near each other in PHATE coordinates. Thus from one end to the next this is akin to a signal condensing and then diffusing again. As expected, this creates a loop-like structure in the PHATE graph. However, directly applying PHATE on the raw signals (**Figure A9J**) results in multiple scattered points separated from the main trajectory, demonstrating that using PHATE only is not able to fully capture and distinguish the signals. Furthermore, although applying PCA on the scattering coefficients (**Figure A9K**) generates two separate trajectories, they fail to form the loop-like structure as with using GSTH. Applying t-SNE (**Figure A9L**) and UMAP (**Figure A9M**) on the generated scattering coefficients also failed to form loop-like structures.

Finally, for the third synthetic dataset, we aim to simulate the propagation of signals similar to that observed in epithelial cells. This will help us to better understand what types of signals GSTH can capture. The propagation of signals among cells can reflect the communication within groups of cells (corresponding to the clusters of Ca^2+^ transients among epithelial cells using Ca^2+^-sensor imaging), while each cell itself can also generate independent signals without communicating with other cells (corresponding to single cells spiking using Ca^2+^ fluorescence imaging). As in Figure A9O for GSTH, the time points formed smooth trajectories at first simulating the propagation of signals on the graph, then disrupted trajectories corresponding to random spiking of single cells. In comparison, using PHATE directly on the raw input signals (**Figure A9Q**) results in more dense clustering (for the initial stage when the signal is just diffusing on the graph) and using t-SNE on the scattering coefficients generates more scattered clustering, making it hard to identify the inner dynamics (**Figure A9S**). Although applying PCA (**Figure A9R**) and UMAP (**Figure A9T**) on the scattering coefficients can reflect part of the dynamics, they also generate very condensed trajectories for the early stage, when the signal is simply diffusing on the graph.

In addition, we computed the Wasserstein distances between the persistence diagram from our GSTH method and persistence diagrams from other methods using the three synthetic datasets (**Figure A9G, A9N, A9U**). The Wasserstein distance between two persistence diagrams *Q*_1_ and *Q*_2_ is calculated as follows:

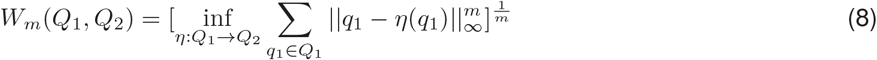

where *η* is a bijection from *Q*_1_ to *Q*_2_. Specifically, we consider the diagonal Δ of persistence diagrams to have infinite multiplicity, i.e. points can be matched to the diagonal. We used Eirene to compute Wasserstein distances, which utilizes the Hungarian algorithm to find the optimal matching. It has been shown that persistence diagrams are stable[41]. Hence by calculating the Wasserstein distances between two persistence diagrams, we can quantify the differences of persistence diagrams. We showed that the persistence diagrams from our GSTH method are different from persistence diagrams produced using other methods ablating different parts of GSTH and visualized the distances with heatmaps (**Figure A9G, A9N, A9U**).

One additional validation of our GSTH approach is based on the observation of similar signaling patterns within experimental groups and consistently different patterns between experimental groups, reflected in the time trajectories. The Wasserstein distances were calculated among persistence diagrams from multiple experimental groups (MMC drug treatment, Cdkn1b overexpression, and control groups) (**Figure 5G**). We found small distances between duplicates from each group and larger distances across different experimental groups, demonstrating GSTH is able to capture homeostatic and perturbed signaling patterns across many samples.

Overall, we show that GSTH can effectively capture signal diffusion dynamics on the graph and is stable to small perturbation of signals. These valuable characteristics of GSTH make it possible to investigate complex spatiotemporal signaling patterns and hence enable us to explore the underlying relationships between Ca^2+^ signaling and specific cell behaviors.

### Cellular Embedding

We can also generate embeddings for individual cell to explore their signaling patterns across time. For each cell, we then consider the signals *X*(*v*_*l*_) = [*X*(*v*_*l*_, *t*_1_),*X*(*v*_*l*_, *t*_2_),…,*X*(*v*_*l*_, *t*_*n*_)], which are defined on cell *v*_*l*_ across all timepoints as features. We utilize the same diffusion operator *R* and graph wavelets **Ψ** defined as in the timepoint embeddings to learn cellular embeddings. Following the calculations in Equation 4, 5 and 6, we can obtain the wavelet coefficients at each vertex/cell. We then concatenate the coefficients of cells across all timepoints to form the cellular embeddings. The cellular embeddings give us description of cell patterns along time, capturing patterns from the cell itself as well as incorporating larger scale signaling patterns by considering neighboring cells at multiple scales.

### Synthetic Dataset for Cell Embedding

Similarly to using synthetic datasets to understand GSTH and the timepoint embeddings, we also created datasets to test our cell embedding methods. Since there are mainly two types of Ca^2+^ signaling patterns observed (single cells spiking and clustered signaling), we aimed to simulate these patterns in the synthetic dataset. Therefore, we created two datasets:

#### Synthetic testcase 4

This dataset contains both types of Ca^2+^ signaling. Specifically, to simulate cells that belong to clustered intercellular Ca^2+^ signaling, we again diffused a Dirac signal *x* defined on node *i* using graph Laplacian *L* for 20 timesteps as:

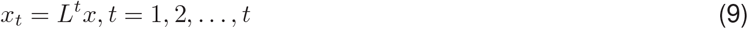

Then for all other cells that do not belong to the Ca^2+^ wave defined above, we defined a single non-diffusing signal on it for a time interval of 5 steps as

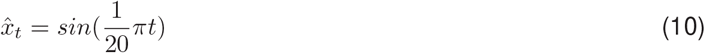

#### Synthetic testcase 5

In the second dataset we consider two intercellular Ca^2+^ waves resulting from two Dirac signals diffused on the graph. The two Dirac signals defined on two different nodes i and j were diffused for 20 timesteps (following Equation 9). This will result in two flashing waves with similar patterns.

### Comparison of Cell Embedding with PHATE using Synthetic Dataset

We first applied geometric scattering to generate wavelet coefficients and then visualized them using PHATE. We colored each data point (each representing a cell in the dataset) with a color scale representing the graph distance to the center cell of each wave, where the Dirac signal was defined initially. We finally compared the results to visualizations where we directly applied PHATE to raw cell signals. For the first synthetic dataset containing both single cell spiking and intercellular Ca^2+^ waves, the geometric scattering transform together with PHATE can clearly distinguish the two types of signaling cells (**Figure A10A, A10C**), with single spiking cells distributed further from the center of flashing waves. In contrast, directly applying PHATE to the cells’ raw signals failed to reveal this pattern, as single spiking cells were condensed, suggesting that PHATE alone is not able to distinguish the graph structure and hence showed less information about graph distances. For the second synthetic dataset containing two similar waves (**Figure A10B, A10D**), cells from the two waves show many overlaps. This is as expected, since although the cells have different spatial locations, their signals are all from the diffusion of Dirac signals on the graph. Thus, they also share similar patterns. Although applying PHATE alone to this dataset also demonstrated similar overlap patterns, the trajectory it generated still failed to reflect the changing graph distance that reflects the graph structure.

## Data and code availability

The source code can be downloaded from https://github.com/krishnaswamylab/GSTH.

## Acknowledgements

We thank David Gonzalez and Kai Mesa for early exploration of calcium signaling in the epidermis and Edward Marsh for help with Python scripts. We also thank all members of the Greco and Krishnaswamy labs for critical feedback on the manuscript. We thank S. Aizawa for the Rosa26p-Fucci2 mice. Finally, we’d like to thank Rachael Norris for her expertise and input on the manuscript.

## Author contributions

J.M., F.G., S.K. and V.G designed experiments. J.M. performed 2-photon imaging, whole mount staining, mouse genetics, and image analysis. F.G. performed data analysis and statistical modeling. C.M.-M. performed 2-photon imaging and image analysis. S.D. and E.L. performed whole mount staining and image analysis.

S.G. performed mouse genetics. L.S. assisted with image analysis. D.B. and B.R. assisted with statistical modeling. J.M., F.G., A.C., C.H., S.K., and V.G wrote the manuscript with input throughout from S.D., C.M.-M., and B.R..

## Data availability

All data from this study are available from the authors on request. The MATLAB and python scripts for the image analysis will be available on request. The source code for GSTH and the cell embeddings can be downloaded from https://github.com/krishnaswamylab/GSTH.

## 1 Appendix

**Figure A1:**
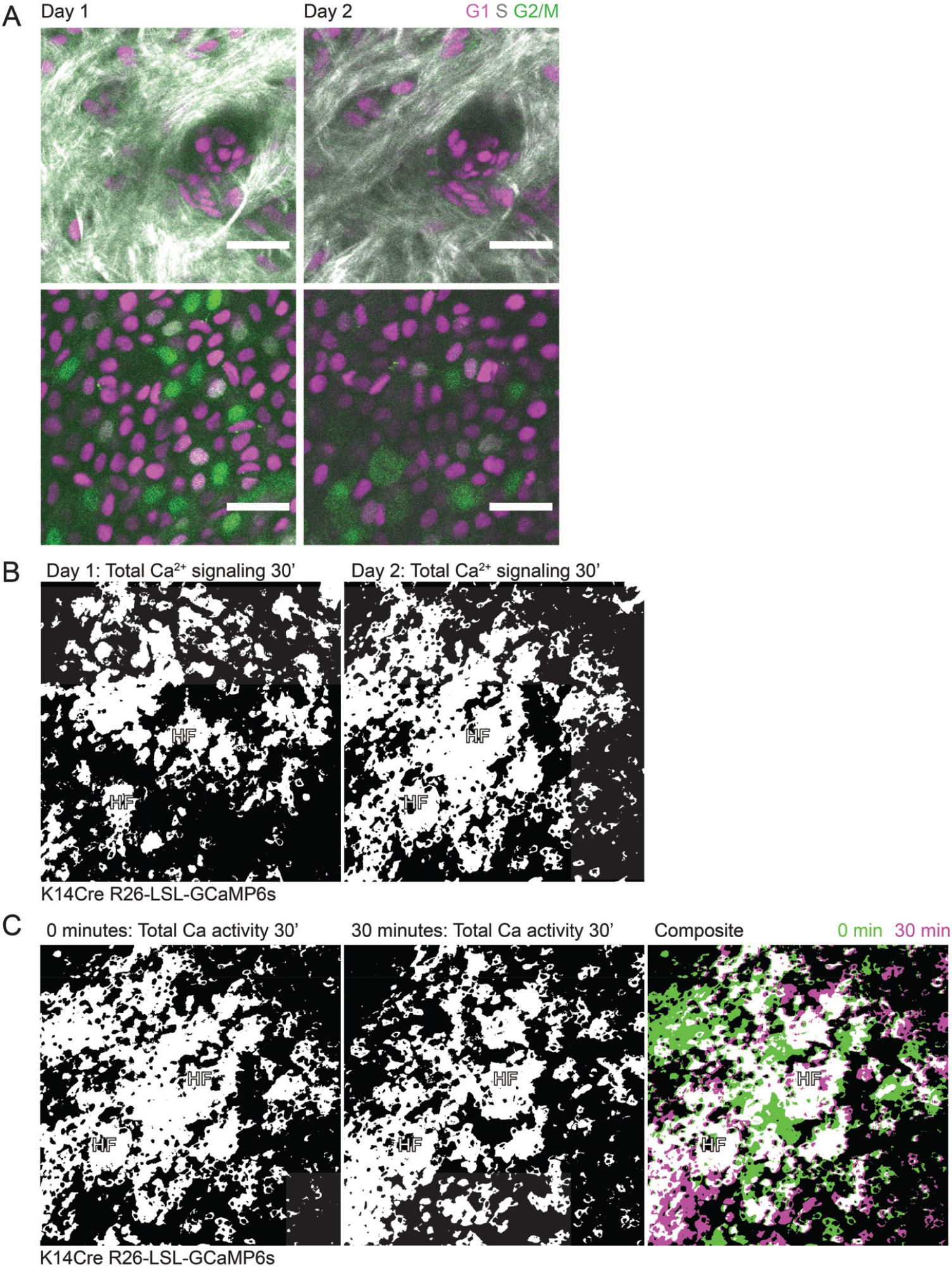
Pervasive, fast Ca^2+^ dynamics are specific to the regenerative basal layer of the epidermis. **(A)** Revisit of the same region of a Rosa26p-Fucci2 mouse at 0 and 24 hours. Top panel show dermis and bottom panel shows epidermal basal layer. Collagen is in white, G1 cells are in magenta, S cells are double positive for magenta and green and shown in gray, G2 and M cells are in green. Scale bars: 25 μm. **(B)** Maximum intensity projection of all optical sections of a 30-minute time-lapse at 0- and 30-minutes of the same region of the epidermis as shown in Figure 1C. Transverse views of the top of the infundibulum region of hair follicles marked with HF to orient us in revisiting the region. **(C)** Maximum intensity projection of all optical sections of a 30-minute time-lapse at 0- and 30-minutes of the same region of the epidermis. To the right, composite image of the same region at 0- (green) and 30-minutes (magenta), where white indicates overlapping regions of Ca^2+^ activity. Transverse views of the top of the infundibulum region of hair follicles marked with HF to orient us in revisiting the region.

**Figure A2:**
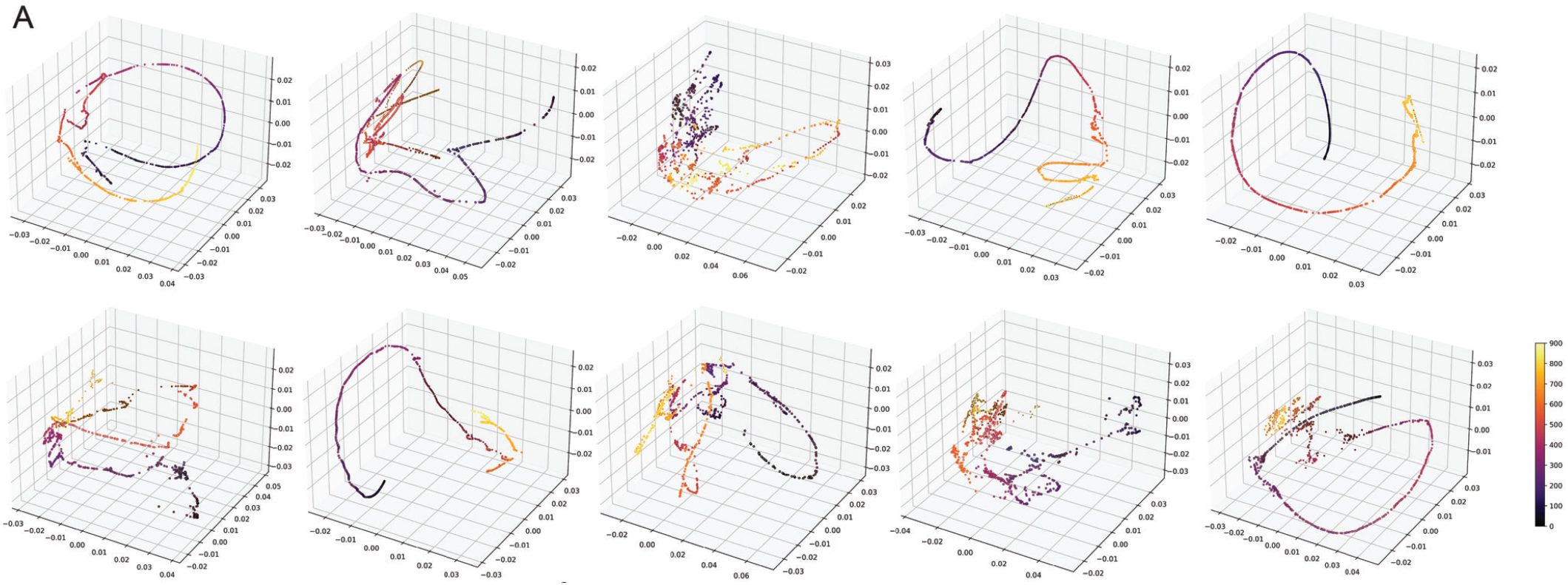
Unsupervised modeling of Ca^2+^ signaling patterns reveals smooth, directed signaling in the homeostatic basal epidermis. **(A)** PHATE visualizations of Ca^2+^ signaling time trajectories in the homeostatic basal epithelial layer from 30-minute time-lapse movies show mainly smooth trajectories.

**Figure A3:**
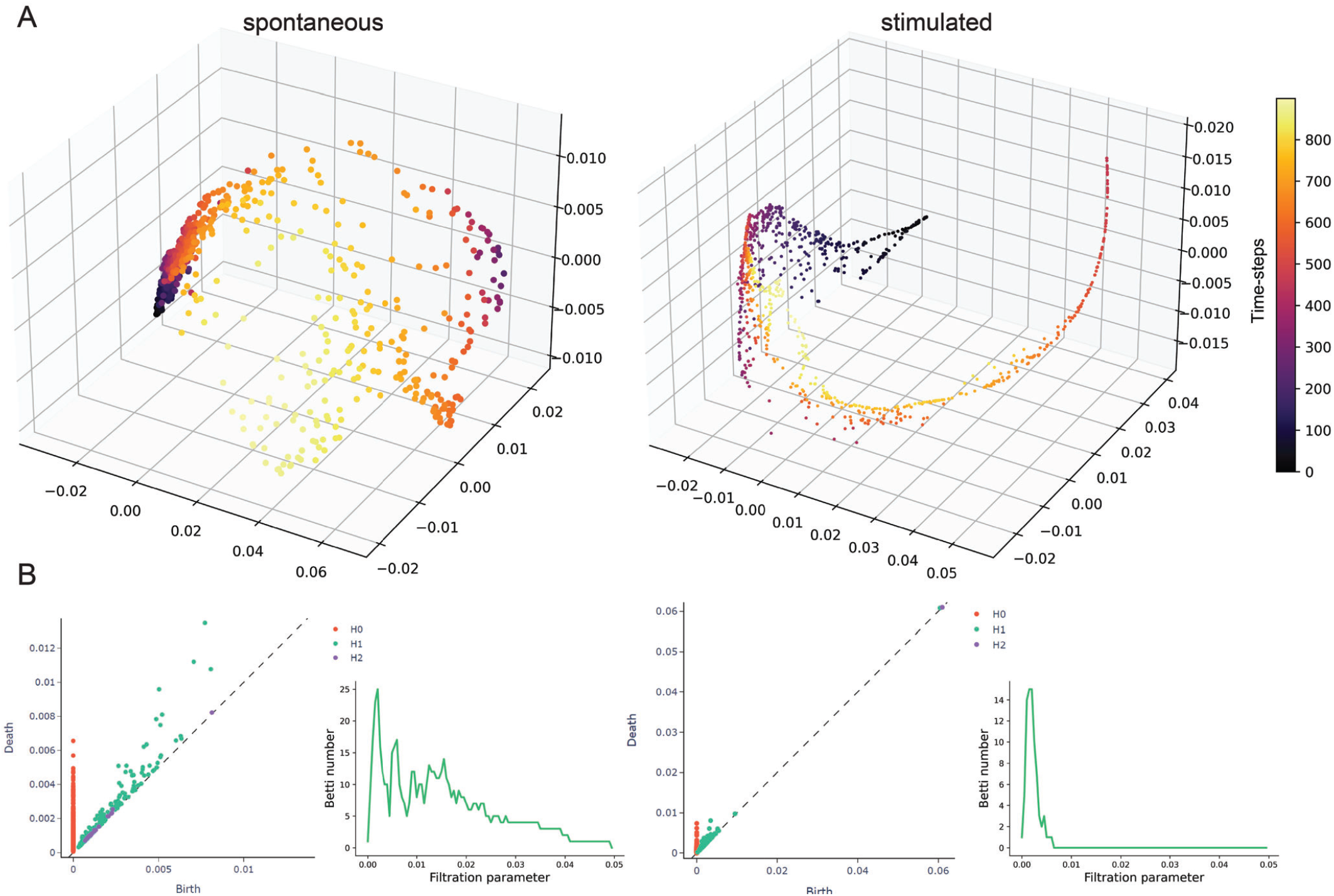
Stimulated visual cortex Ca^2+^ signaling is more temporally coordinated than spontaneous activity. **(A)** PHATE visualization of Ca^2+^ signaling patterns in visual cortex with spontaneous or stimulated neuronal activity. **(B)** Representative persistence diagrams (*H*0: connected components, *H*1: loops, *H*2: voids) and Betti curves of *H*1 features for spontaneous and stimulated neurons of the visual cortex. The persistence diagram for spontaneous activity has a rich collection of *H*2 features/voids, which is less common in stimulated activity. In addition, the *H*1 feature/loops from the spontaneous activity shows longer persistence

**Figure A4:**
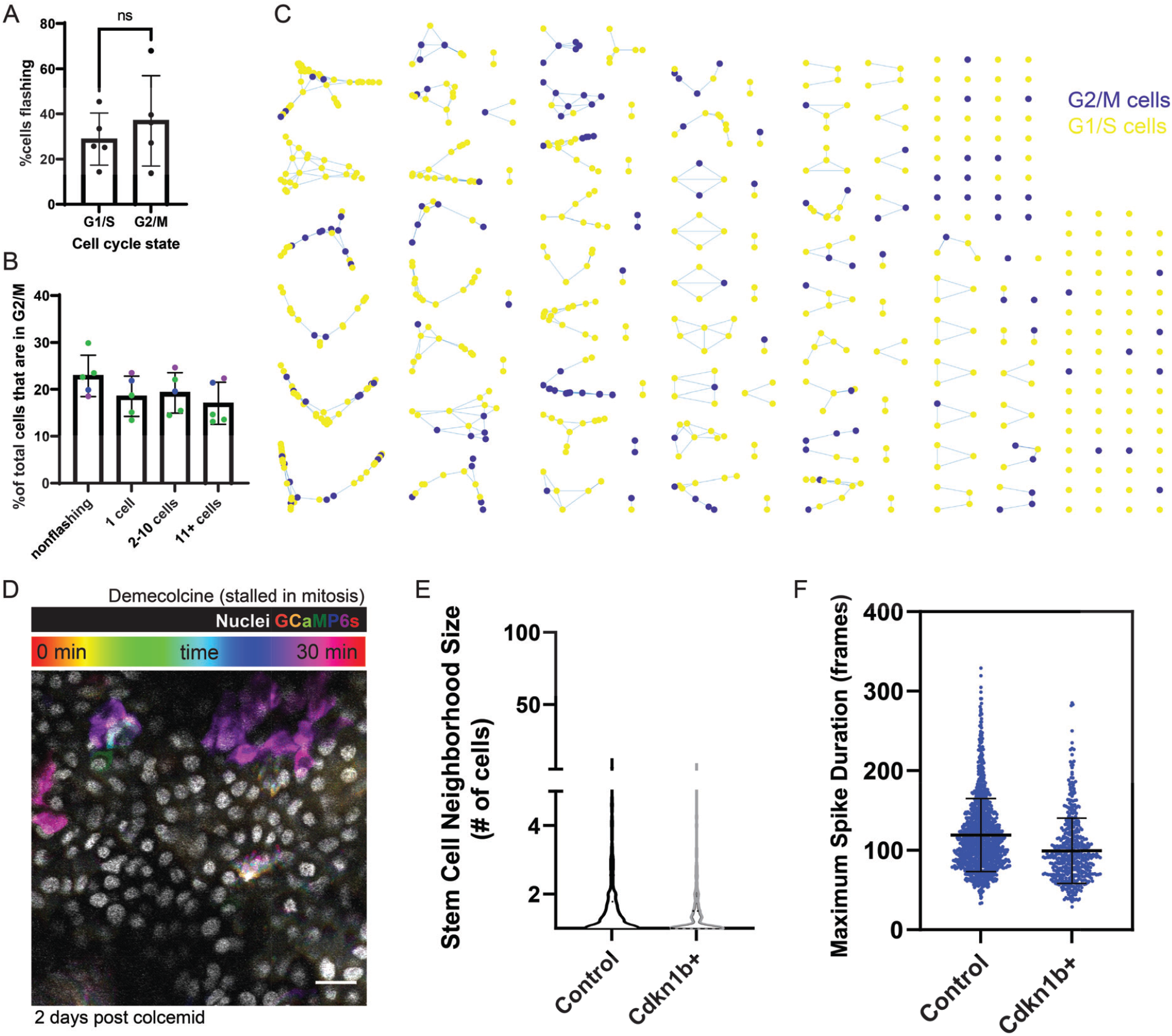
Cell cycle specific Ca^2+^ signaling. **(A)** Maximum intensity projection of a 30-minute time-lapse video of the epidermal basal layer of a live Ca^2+^ reporter mouse two days after treatment with demecolcine stalling cells in mitosis (K14-Cre; Rosa26-CAG-LSL-GCaMP6s; K14-H2BmCherry). Ca^2+^ sensor fluorescence over time is represented as a color scale and nuclei are shown in white. Scale bar: 25 μm. **(B)** Percent of G1/S versus G2/M cells flashing over the course of 30 minutes. N = 5 thirty-minute time-lapse movies from 3 individual mice. **(C)** Percent of G2/M cells in groups of non-flashing, single flashing, small clusters, and large clusters of flashing cells based on mCherry-hCdt1 expression. N = 5 thirty-minute time-lapse movies from 3 individual mice. **(D)** “Neighborhoods” of spatiotemporally connected Ca^2+^ signaling colored by cell cycle stage (blue = G1/S and yellow = G2/M) from a representative 30-minute time-lapse movie. **(E)** Histogram showing relative frequency of different neighborhood sizes of spatiotemporally connected Ca^2+^ signaling from 30-minute time-lapses of cdkn1b+ G1-stalled basal layers (green) versus control (blue) in Ca^2+^ sensor mice. **(F)** Maximal spike duration (maximum number of frames between the start and end of individual Ca^2+^ events) in control versus G1-stalled cdkn1b+ mice. Bars denote mean and error bars represent SD. N = 9 control and 8 cdkn1b+ thirty-minute time-lapse movies from at least 3 mice per condition.

**Figure A5:**
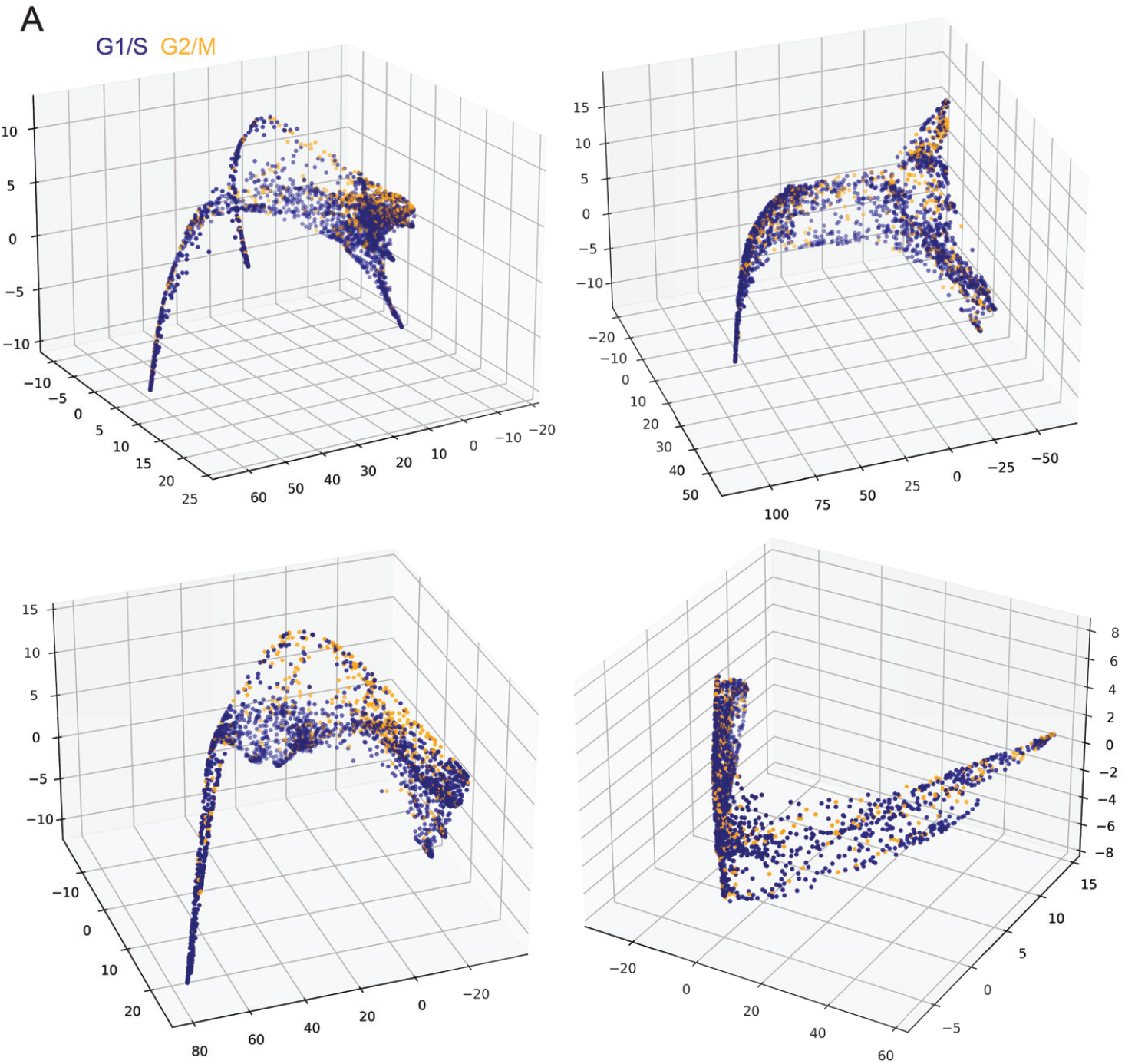
G2 cells’ Ca^2+^ traces cluster more closely than G1 or S cells’ Ca^2+^ traces. **(A)** PHATE visualization of cell clustering of Ca^2+^ signaling patterns, where each dot represents a single cell; its position in space represents how similar its Ca^2+^ signaling is to other cells in space; each cell or node is colored by its cell cycle state based on nuclear Fucci2 signal.

**Figure A6:**
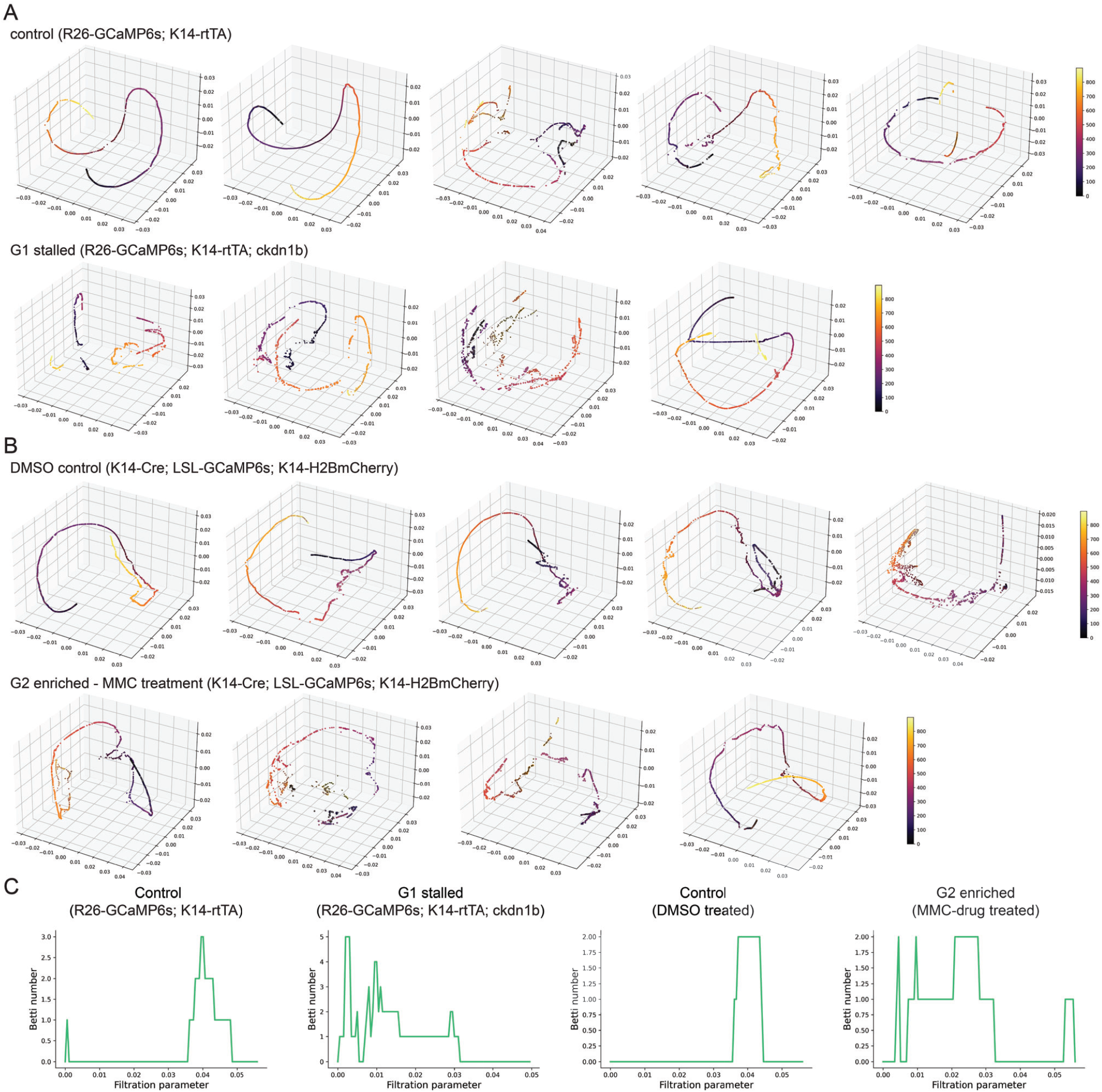
Enrichment of G2 cells uniquely does not disrupt coordinated Ca^2+^ signaling. **(A)** PHATE visualization of Ca^2+^ signaling in the cdkn1b+ G1-stalled basal layer versus control shows disruption of smooth, directed and coordinated patterns of signaling. **(B)** PHATE visualization of Ca^2+^ signaling in the MMC-treated G2-enriched basal layer versus control shows smooth, directed and coordinated patterns of signaling. **(C)** Representative Betti curves of *H*1 features (loops) for G1 and G2 enriched conditions (R26-GCaMP6s; K14-rtTA; cdkn1b 3 days after doxycycline treatment and MMC drug 2 days after treatment respectively)

**Figure A7:**
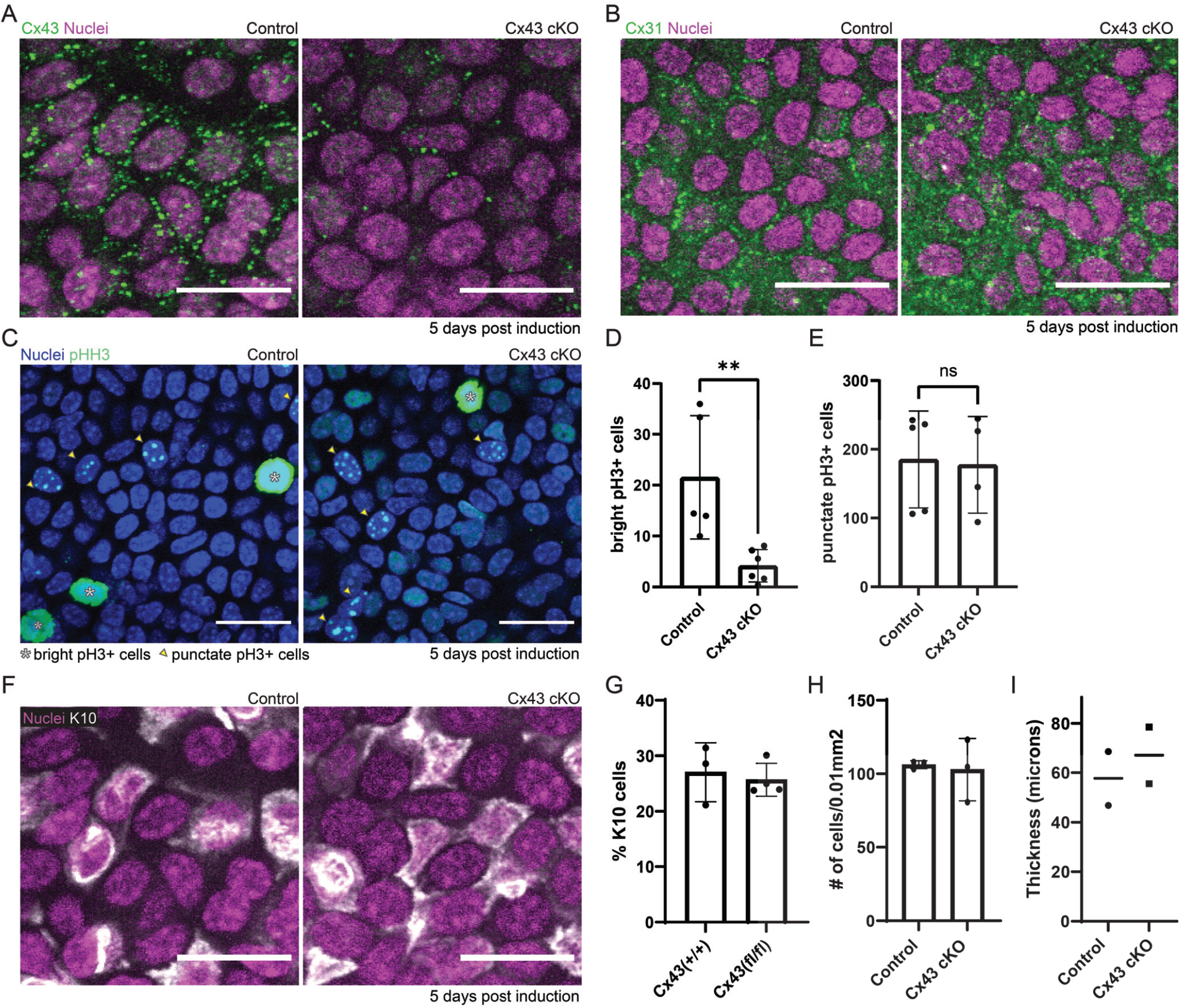
Loss of Cx43 does not fully disrupt all gap junctions or the integrity of the tissue. **(A)** Immunofluorescence staining of basal layer from K14-CreER; Cx43^+/+^ and K14-CreER; Cx43^fl/fl^ mice 5 days post-tamoxifen induction. Cx43 in green and DAPI (nuclei) in magenta. Scale bar: 25 μm. **(B)** Immunofluorescence staining of epidermal basal stem cell layer from K14-CreER; Cx43^+/+^ and K14-CreER; Cx43^fl/fl^ mice 5 days post tamoxifen induction, with staining for Cx31 in green and Hoechst marking nuclei in magenta. Scale bar: 25 μm. **(C)** Phospho-histone H3 (pH3) immunofluorescence staining in control versus Cx43 cKO mice (K14-CreER Cx43^+/+^ and K14-CreER Cx43^fl/fl^) 5 days post-tamoxifen induction. Scale bars: 25 μm. **(D)** Quantification of number of bright pH3 positive cells per 250 mm^2^ region 5 days post-tamoxifen induction, marking mitotic cells. ** P *<*0.01, Student’s t test. N = 5 control and 6 Cx43 cKO mice. **(E)** Quantification of punctate pH3 positive cells per 250 mm^2^ region in control versus Cx43 cKO mice 5 days post-tamoxifen induction, marking late G2 cells. N = 5 control and 4 Cx43 cKO mice. **(F)** K10 immunofluorescence staining in control versus Cx43 cKO mice (K14-CreER Cx43^+/+^ and K14-CreER Cx43^fl/fl^ respectively) 5 days post-tamoxifen induction. Scale bars: 25 μm. **(G)** Quantification of K10 positive basal cells as a percentage of total basal cells 5 days post-tamoxifen induction in Cx43 cKO and control mice, indicating cells that are beginning differentiation. NS, Student’s t test. N = four 10 mm^2^ regions per mouse, 3 mice per experimental group. **(H)** Quantification of average cell density in control versus Cx43 cKO mice (K14-CreER Cx43^+/+^ and K14-CreER Cx43^fl/fl^ respectively) 5 days post-tamoxifen induction. N = six 10 mm^2^ regions per mouse, 3 mice per experimental group. **(I)** Quantification of epidermal thickness in control versus Cx43 cKO mice (K14-CreER Cx43^+/+^ and K14-CreER Cx43^fl/fl^ respectively) 5 days post-tamoxifen induction. N = six 10 mm^2^ regions per mouse, 2 mice per experimental group.

**Figure A8:**
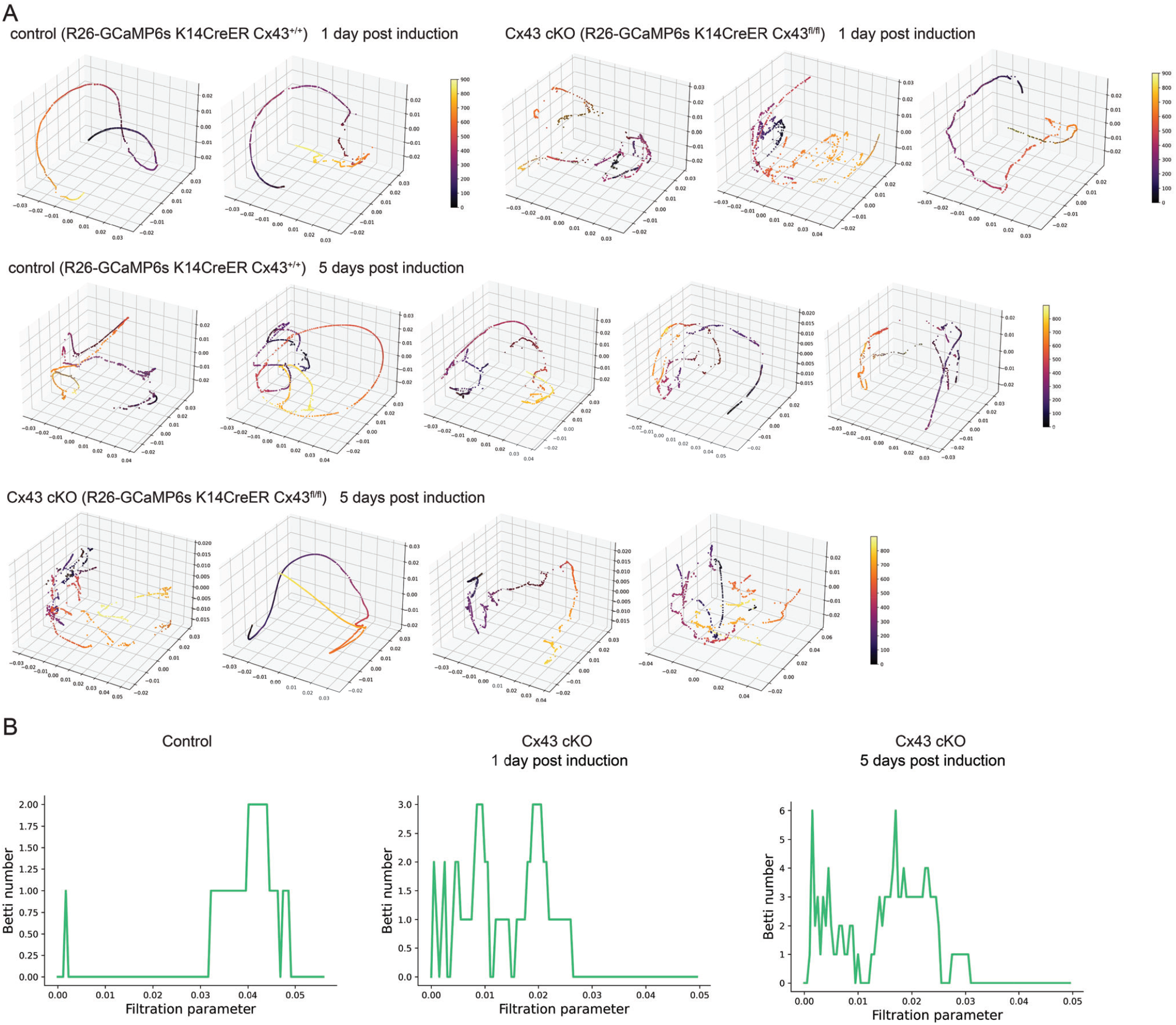
Loss of Cx43 disrupts coordinated Ca^2+^ signaling patterns. **(A)** PHATE visualization of Ca^2+^ signaling time trajectories in the Cx43 conditional knockout versus control basal layer shows disruption of smooth, directed and coordinated patterns of signaling in mice 1 and 5 days after loss of Cx43. **(B)** Representative Betti curves of *H*1 features (loops) for control and Cx43 cKO mice (Rosa26-CAG-GCaMP6s; K14-CreER; Cx43^+/+^ and Rosa26-CAG-GCaMP6s; K14-CreER; Cx43^fl/fl^) 1 and 5 days post-tamoxifen induction.

**Figure A9:**
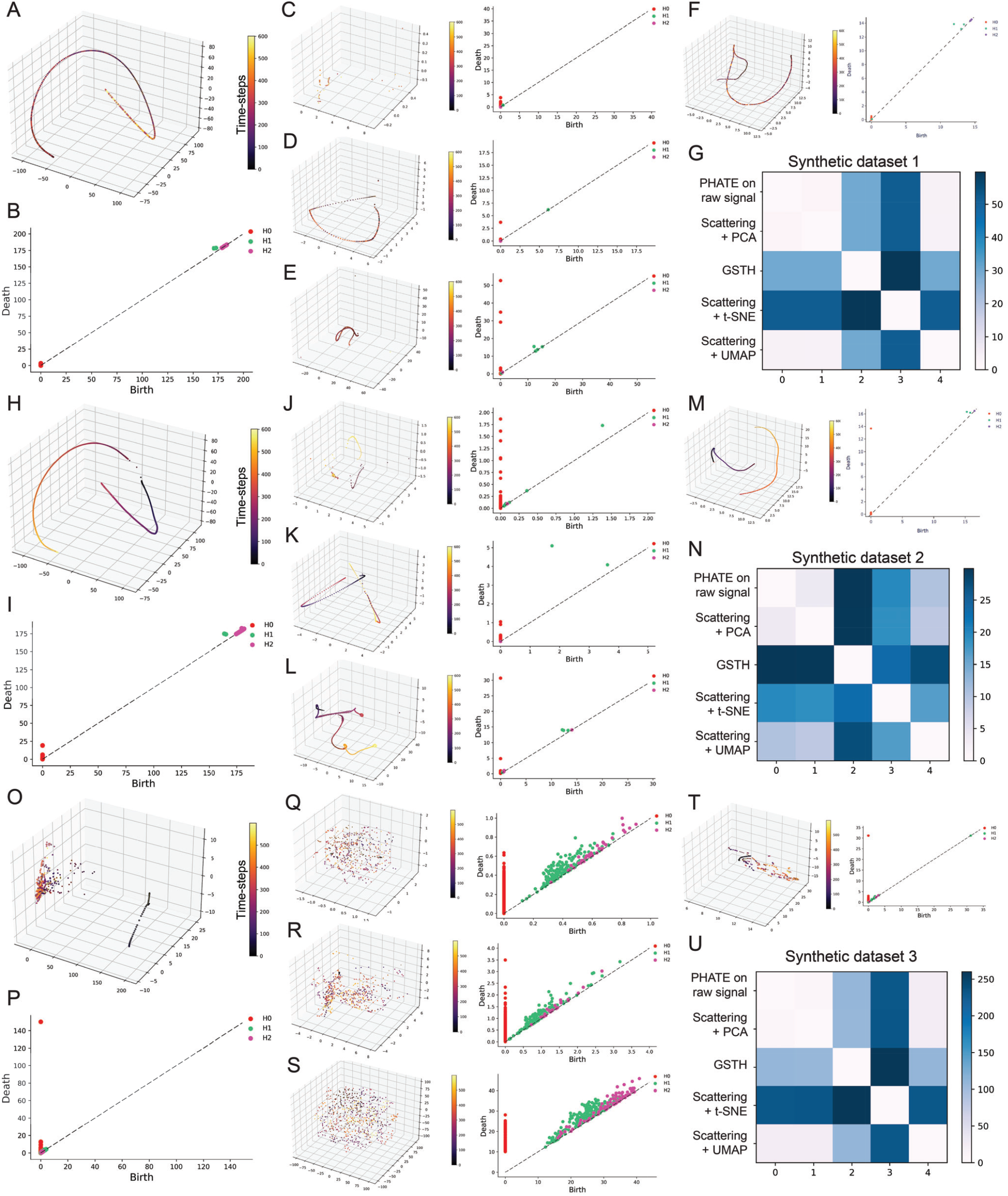
Validation of GSTH using synthetic datasets. PHATE visualization **(A, H, O)** and persistence homology **(B, I, P)** on synthetic data using GSTH and comparison with (1) directly applying PHATE on the input signals **(C, J, Q)**; (2) PCA on generated scattering coefficients **(D, K, R)**; (3) t-SNE on generated scattering coefficients **(E, L, S)**; (4) UMAP on generated scattering coefficients **(F, M, T)**. Finally, Wasserstein distances from the persistence diagrams of each methodology for each of the 3 synthetic datasets **(G, N, and U)**

**Figure A10:**
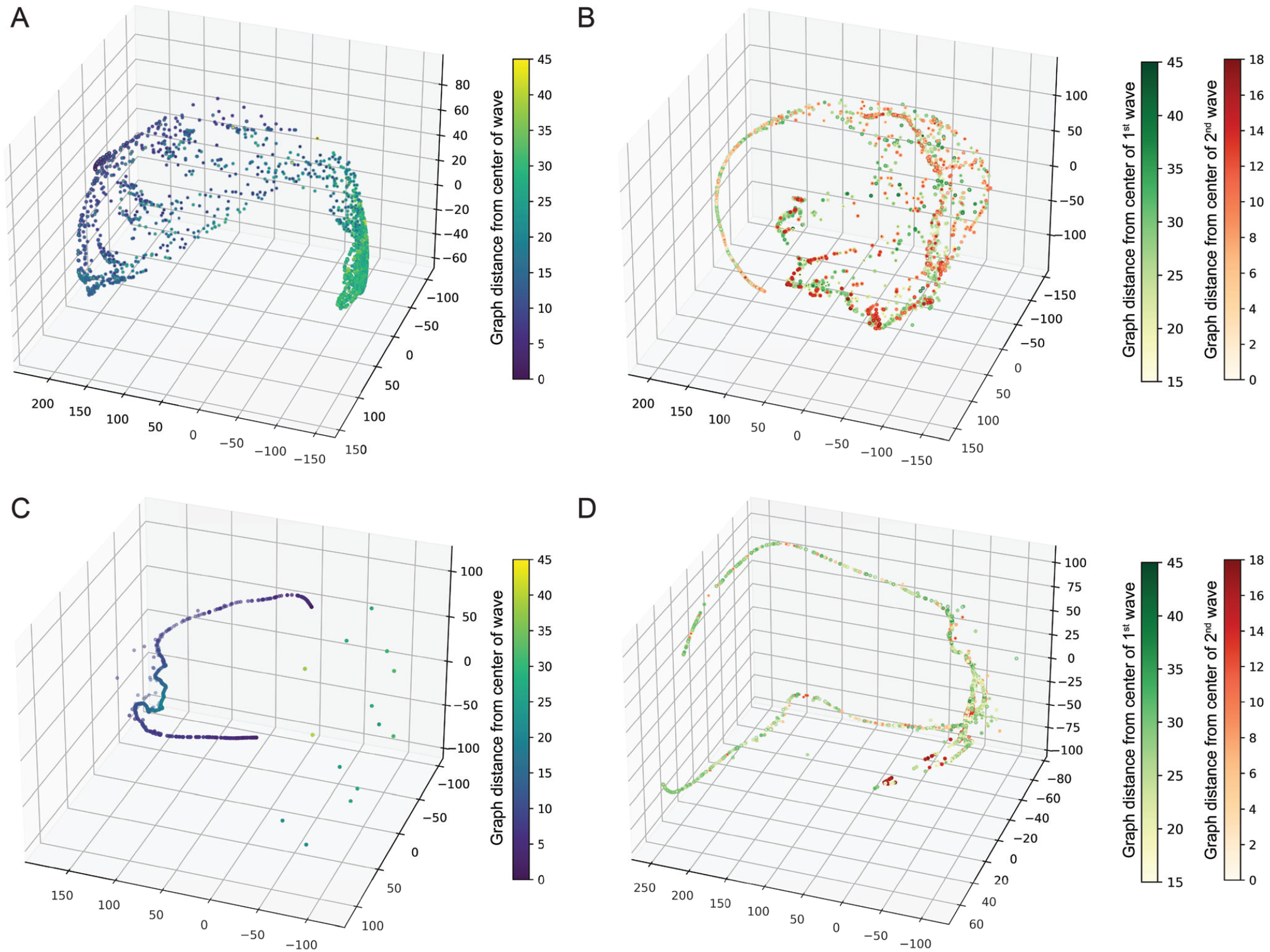
Validation of cellular embeddings using synthetic datasets. **(A)** PHATE visualization of cellular embeddings generated with geometric scattering transform for synthetic dataset 4. **(B)** PHATE visualization of cellular embeddings generated with geometric scattering transform for synthetic dataset 5. **(C)** PHATE visualization alone of cells with raw signals for synthetic dataset 4. **(D)** PHATE visualization alone of cells with raw signals for synthetic dataset 5.

