## Supplementary material for "G2 stem cells orchestrate time-directed, long-range coordination of calcium signaling during skin epidermal regeneration": Movie Legends

**Movie 1. Variable participation in $Ca^{2+}$ signaling across the basal stem cell layer.** Intravital imaging of the epithelial stem cell layer of $Ca^{2+}$ sensor mice (K14-Cre; R26-LSL-GCaMP6s; K14-H2BmCherry). Magenta marks the nuclei of each basal cell and green fluorescence intensity represents relative cytosolic $Ca^{2+}$ levels. Transverse views of the infundibulum of hair follicles marked with HF. Frames taken every 2 seconds for a total duration of 30 minutes. Movie shows dynamics at 40 frames per second or 80x the actual timescale. Scale bar: 25 µm.

**Movie 2. Mitotic cells do not participate in homeostatic $Ca^{2+}$ signaling.** Intravital imaging of $Ca^{2+}$ sensor mice (K14-Cre; R26-LSL-GCaMP6s; K14-H2BmCherry) with eight examples of basal epithelial stem cells undergoing mitosis showing a lack of $Ca^{2+}$ signaling. Magenta marks the nuclei of each stem cell and green fluorescence intensity represents relative cytosolic $Ca^{2+}$ levels. Frames taken every 2 seconds for a total duration of 30 minutes. Movie shows dynamics at 60 frames per second or 120x the actual timescale. Scale bar: 10 µm.

**Movie 3. Basal stem cells stalled in mitosis show low levels of $Ca^{2+}$ signaling.** Disrupted $Ca^{2+}$ dynamics in demecolcine-treated mice on the right compared to DMSO vehicle-treated littermate controls on the left, 2 days after drug treatment. Magenta marks the nuclei of each stem cell and green fluorescence intensity represents relative cytosolic $Ca^{2+}$ levels. Frames taken every 2 seconds for a total duration of 23 minutes. Movie shows dynamics at 40 frames per second or 80x the actual timescale. Scale bar: 25 µm.

**Movie 4. Basal stem cells stalled in G1 of their cell cycle show low levels of $Ca^{2+}$ signaling.** Disrupted $Ca^{2+}$ dynamics in G1-stalled Cdkn1b mice (K14-rtTA; tetO-Cdkn1b; R26-GCaMP6s) on the right compared to littermate controls (K14-rtTA; R26-GCaMP6s) on the left, 3 days after start of doxycycline administration. Green fluorescence intensity represents relative cytosolic $Ca^{2+}$ levels. Frames taken every 2 seconds for a total duration of 30 minutes. Movie shows dynamics at 40 frames per second or 80x the actual timescale. Scale bar: 25 µm.

**Movie 5. Basal stem cells stalled in G2 of their cell cycle show normal patterns of $Ca^{2+}$ signaling.** G2-enriched Mitomycin C-treated mice on the right compared to DMSO vehicle-treated littermate controls on the left, 2 days after drug treatment. Magenta marks the nuclei of each basal cell and green fluorescence intensity represents relative cytosolic $Ca^{2+}$ levels. Frames taken every 2 seconds for a total duration of 30 minutes. Movie shows dynamics at 40 frames per second or 80x the actual timescale. Scale bar: 25 µm.

**Movie 6. Cx43 orchestrates $Ca^{2+}$ signaling at large scales, but not across local neighborhoods, in the stem cell pool.** Disrupted $Ca^{2+}$ dynamics in Cx43 cKO mice (K14CreER; $Cx43^{fl/fl}$; R26-GCaMP6s) on the right compared to littermate controls (K14CreER; $Cx43^{+/+}$; R26-GCaMP6s) on the left, 5 days after tamoxifen induction. Green fluorescence intensity represents relative cytosolic $Ca^{2+}$ levels. Frames taken every 2 seconds for a total duration of 30 minutes. Movie shows dynamics at 40 frames per second or 80x the actual timescale. Scale bar: 25 µm.
